# Whole-genome resequencing and indoor phenotyping resolve the genetic architecture of drought stress response in tepary bean

**DOI:** 10.64898/2026.11.24.768977

**Authors:** Aparna Srinivasan, Li’ang Yu, Ruth Nichols, Emily Miller, Annie Voss, Andrew D. L. Nelson, Duke Pauli, Timothy Porch, Magdalena M. Julkowska

## Abstract

Tepary bean (*Phaseolus acutifolius* A. Gray) is an arid-adapted legume, whose drought resilience is widely reported but poorly resolved genetically. Here we resequenced 363 wild, weedy, cultivated and breeding accessions and recovered 6.9 million single-nucleotide polymorphisms (SNPs). We phenotyped 282 of the accessions for growth, evapotranspiration and photosynthesis over 14 days at 60% or 10% soil water-holding capacity. Cultivated and breeding lines grew roughly twice as fast as wild accessions, and the proportional reduction in growth under drought did not differ between these germplasm classes. Evapotranspiration separated the classes under full irrigation, but converged under drought, making it the most plastic trait we measured. Canopy density predicted how much water a plant used, but not how well it tolerated losing it. Association mapping across 104 rank-normalized measurements yielded 113 significant SNPs at 46 positions, which resolved to three drought and six control loci. Three drought loci corresponded to 98 of the 113 SNPs, with the largest one localized on chromosome 8. This locus separates wild from cultivated accessions, and resolves to no gene. Three of the eight remaining loci carry candidates whose interval, expression under drought and published mechanism agree: a cytokinin oxidase, a CIPK1-clade kinase and the antenna subunit LHCB5. The most structure-independent locus localizes to 26 kb but implicates genes with no published role in water relations. Growth under drought and control conditions therefore rest on separate genetic bases, and the variation with the largest effect under deficit lies in haplotypes affected by domestication.

**Highlight:** Domestication doubled tepary bean vigor without eroding drought resilience. Correcting for plant size and for population structure reduces 113 significant SNPs to nine loci, three of which carry candidate genes.

## Introduction

Domestication and breeding typically increase yield, but they can also reduce genetic variation. That reduction is widely understood to limit the capacity of modern cultivars to tolerate abiotic stress (Warschefsky *et al*., 2014; von Wettberg *et al*., 2018). Modern agriculture depends on a handful of species (Massawe *et al*., 2016), while the environmental resilience held in indigenous crops and landraces remains underutilized (Dwivedi *et al*., 2016). Therefore, broadening the resilience available to breeders requires identifying allelic variation in dryland agriculture species.

Tepary bean (*Phaseolus acutifolius* A. Gray) is a legume native to arid and semi-arid regions of the southwestern United States, northwestern Mexico and Central America (Nabhan and Felger, 1978; Garvin and Weeden, 1994). It is traditionally sown during two planting seasons in the Sonoran Desert, and its nutritional profile is comparable to common bean but higher in sucrose and lower in fat (Porch *et al*., 2017). Its stress resilience has been recognized repeatedly (Markhart, 1985; Federici *et al*., 1990; Rainey and Griffiths, 2004; Rao *et al*., 2013; Buitrago-Bitar *et al*., 2021). Recent work documents substantial variation in stomatal and hydraulic drought responses within and between *Phaseolus* species (Buckley *et al*., 2025). Breeders have introgressed tepary alleles into common bean to transfer abiotic stress tolerance (Souter *et al*., 2017), and tepary-focused breeding has raised yields (Porch *et al*., 2017; reviewed in Mhlaba *et al*., 2018 and Mwale *et al*., 2020). The species nonetheless remains largely absent from molecular studies of stress biology.

Recently, tepary bean gained its genomic foundation in the form of a reference genome (Moghaddam *et al*., 2021) and two diversity panels. The first panel consisted of 314 accessions evaluated by genotyping-by-sequencing and resequencing (Moghaddam *et al*., 2021). The second panel phenotyped 422 accessions for biotic stress resistance, resolving in eight subpopulations, and mapped loci for weevil, common bacterial blight, *Fusarium* wilt and BCMNV resistance (Bornowski *et al*., 2023). Both confirmed that domestication compressed diversity, consistent with a single domestication event (Garvin and Weeden, 1994; Gujaria-Verma *et al*., 2016). Their SNP densities of 20,364 and 53,676 SNPs are too sparse to resolve individual loci for quantitative traits, and neither was phenotyped for drought.

Here we re-sequenced 363 tepary accessions, resulting in 6.9 million SNPs, and phenotyped 282 accessions under a sustained water deficit. To separate the response to water limitation from the developmental differences between the accessions, the plants were grown in a controlled environment, watered to 60 and 10% of soil water-holding capacity and phenotyped in detail over 2 weeks of experimentation. The collected data was examined across three axes. First, we examined whether domestication and breeding reduced drought resilience in favor of seedling vigor. Second, we explored which measured traits predicted the maintenance of growth under water limitation once plant size is accounted for. Third, we identified regions of the genome corresponding to variation in both development and drought stress response. All three exploratory axes converged, suggesting that performance under water deficit is not a scaled-down version of performance under sufficient irrigation, but rather a separate target with its own predictive traits and genetic loci.

## Materials and methods

### Plant genetic material

In total, 363 *Phaseolus acutifolius* accessions (**Table S1**) were used in this study. One generation of single-seed descent was used to purify the Tepary Diversity Panel at the USDA-ARS-TARS in Mayagüez, Puerto Rico. The parental generation used to produce seed for this study was tested for the presence of *Bean common mosaic virus* (BCMV) and *Bean common mosaic necrosis virus* (BCMNV) with an Agdia (Elkhart, IN, USA) ELISA kit. Only accessions without detectable BCMV or BCMNV were used for further experiments.

### DNA extraction

Young, actively growing leaf tissue was harvested, and DNA was isolated using the Wizard® Genomic DNA Purification Kit (Promega Corporation, Madison, WI, USA) following the manufacturer’s protocol. The quality of the DNA was evaluated using a 1% agarose gel. Samples that exhibited high RNA contamination were treated with 10 mg/ml RNase A, incubated at 37 °C for one hour, and purified using the Macherey-Nagel™ NucleoSpin™ gDNA Clean-up Kit (MACHEREY-NAGEL GmbH & Co. KG, Düren, Germany). The DNA samples were sent to Novogene Corporation Inc. (Sacramento, CA, USA) for whole-genome sequencing, which included quality control, preparation of a 350 bp insert DNA library, and sequencing using Novaseq PE150.

### Sequencing reads processing and variants calling

Per-accession read, mapping and variant-calling statistics are given in **Tables S2** and **S3**. The Illumina reads of all tepary accessions (PE 150 bp) were processed by removing adaptors and reads with low quality using fastp (Chen *et al*., 2018; parameters: –cut_tail_mean_quality 15, – n_base_limit 5, –average_qual 10). The cleaned reads were mapped to the domesticated tepary bean reference genome (G40001, *P. acutifolius*, Moghaddam *et al*., 2021) using BWA-mem (v0.7.17-r1188; Li, 2013) with default parameters. Duplicated read-pairs and low-quality mapping in each sample were cleaned using sambamba (Tarasov *et al*., 2015) v1.28.1, parameter: mapping_quality >= 1 and not (unmapped or secondary_alignment) and not (] != null or [SA] != null), followed by read sorting in samtools (Danecek *et al*., 2021). Variants were then called using the GATK4 “HaplotypeCaller” function (v4.2.5.0; McKenna *et al*., 2010, parameters: –max-alternate-alleles 6, –Ploidy 2). The single nucleotide polymorphisms (SNPs) were filtered using the GATK hard-filtering protocol ‘QD < 2.0 || FS > 60.0 || MQ < 40.0 || MQRankSum < 12.5 || ReadPosRankSum < 8.0’, and insertions and deletions (INDELs) with parameters ‘QD < 2.0 || FS > 200.0 || ReadPosRankSum < 20.0’. Those high-quality SNPs and INDELs were further annotated by the variant effect predictor (VEP, McLaren *et al*., 2016) using the *P. acutifolius* gene annotation (Moghaddam *et al*., 2021).

### Population structure analyses

For SNPs retained from GATK variant calling, only bi-allelic SNPs at the four-fold degenerate (4DTV) loci were filtered by plink (v1.90b4, –types SNPs -m 2 -M 2) for population structure analysis. The called set was pruned for linkage disequilibrium (LD) in plink (Purcell *et al*., 2007; –indep-pairwise 200 50 0.2, MAF ≥ 0.05), leaving 95,311 SNPs, which were used for principal component analysis in GCTA (Yang *et al*., 2011). Removing SNPs missing in more than 10% of accessions left 65,596, on which ADMIXTURE (Alexander, Novembre, and Lange, 2009) was run for *K* = 2 to 10 with ten-fold cross-validation (**Fig. S1**).

### Characterization of drought stress-induced changes under controlled conditions

A total of 282 accessions were screened in the experiment (**Table S4**), including 144 wild, 106 cultivated, 8 weedy, and 24 breeding lines. The accessions are drawn from the Tepary Diversity Panel assembled by Bornowski *et al*. (2023) from the CIAT and USDA National Plant Germplasm System collections, and retain its TDP identifiers: 259 of the 282 phenotyped accessions (92%) are TDP members, covering 61% of that 422-accession panel, and 350 of the 363 sequenced accessions carry TDP identifiers; the remaining 13 are the previously published accessions of Moghaddam *et al*. (2021). The subset is enriched for wild germplasm relative to the full panel — wild and weedy accessions make up 55.6% of those phenotyped here against 46.0% of the TDP, and the wild variety *tenuifolius* 32% against 24% — because the large, predominantly cultivated subpopulation of that panel is sampled at 42% while its wild subpopulations are sampled at 77 to 83% (**Table S11**). Accession-level correspondence is given in **Table S11**. Plants were grown in a walk-in growth chamber with a 16 h light / 8 h dark cycle at 25 °C during the day and 21 °C at night, and 60% humidity. Seeds were subjected to scarification prior to germination on germination paper. Upon the development of a radicle, seedlings were transplanted into 10 cm pots filled with a growth medium comprised of Cornell Mix (0.16 m3 of peat moss, 20.84 kg of vermiculite, 0.59 kg of Uni-Mix fertilizer, and 2.2 kg of lime) plus Osmocote (ICL Group, St. Louis, MO, USA). The accessions were distributed across eight experimental batches, with two accessions serving as checks in all experiments: TDP-22 (cultivated) and TDP-359 (wild). Control plants were maintained at 60% soil water-holding capacity (SWHC), while drought-treated plants were kept at 10% SWHC. To determine the target weights for each pot, a drying period of 72 hours was employed, with this weight designated as indicative of 0% soil water-holding capacity. Subsequently, the soil was fully saturated for 24 hours, excess water was drained, and the pot weights were measured, establishing this value as 100% soil water-holding capacity. Monitoring of pot weights commenced 14 days after germination on the AAWESMO gravimetric platform (Yu *et al*., 2024), which weighed each pot daily and watered it back to its predetermined target weight. Daily evapotranspiration was recorded as the weight lost by each pot over the preceding 24 h. The target weights for the drought treatment were reached on the third day of monitoring. Starting from 14 days post-germination, plant imaging was conducted using the PhenoCage system (Yu *et al*., 2024) every two days for the subsequent two weeks, resulting in eight time-points, each comprising seven images per plant. Digital biomass was then interpolated to a daily series by the smooth-spline fit described below, so that even-numbered days are measured and odd-numbered days are interpolated; of the five days carried into the association analysis, days 10, 12 and 14 are measured and days 11 and 13 interpolated. Evapotranspiration, by contrast, was measured every day. Photosynthetic capacity was measured with a PhotoSynQ device on days 6 and 13 of the imaging series, which is 4 and 11 days after the drought pots reached their target weights; days are numbered from the start of imaging throughout this paper. Measurements were made on light-adapted leaves; plants were not dark-adapted, so the maximum quantum yield of photosystem II (Fv/Fm) was not measured and is not reported. The parameters retained were the minimum and maximum fluorescence of a light-adapted leaf (Fo′, Fm′), the operating efficiency of photosystem II (Fv′/Fm′), the quantum yields of photosystem II photochemistry, of regulated and of unregulated non-photochemical quenching (ϕII, ϕNPQ, ϕNO), non-photochemical quenching (NPQ(t)), the fraction of open photosystem II centers (qL), linear electron flow, the SPAD chlorophyll index, leaf temperature and leaf thickness. The photosystem I active- and open-centre parameters returned by the device were discarded: both are defined on the interval zero to one, and 73% and 61% of their values respectively fell outside it in the raw instrument output, so the P700 measurement is taken to have failed in this setup. The three quantum yields sum to unity in every record, which is the internal consistency check for the fluorescence measurements that were retained. At the end of the experiment (15 days after stress imposition), fresh and dry shoot weight was collected for all imaged plants.

### Phenotypic data curation for plants grown under controlled conditions

The collected images of tepary beans were processed using OpenCV (Bradski 2000) and the PlantCV library (Gehan *et al*., 2017). The notebook describing the full pipeline, consisting of (1) extracting unique pot names from image filenames, (2) mapping unique pots to their respective ArUco IDs, and (3) analyzing images to obtain plant area, height, and width, can be accessed at https://doi.org/10.5061/dryad.9w0vt4bwb. The collected raw data can be accessed at https://doi.org/10.5061/dryad.9w0vt4bwb. All results were processed in R; the image-processing pipeline is available at https://github.com/aparnahorti/BTI_Tepary_Bean_Phenotyping_Pipeline. Individual experimental datasets were combined, and the increase in shoot projected area and evapotranspiration were modeled using smooth splines. Growth rate is the slope of a linear fit to the digital biomass series of each plant. Cumulative evapotranspiration (EVT.sum) is the sum of the daily values over the treatment, and median evapotranspiration (EVT.med) is the median of days 4 to 11. Water content is fresh weight minus dry weight, and evapotranspiration per unit fresh weight is cumulative evapotranspiration divided by harvest fresh weight. Daily evapotranspiration values that were negative or physiologically impossible were replaced by the spline fit to the remaining days for that plant. The corrected phenotypic data (**Table S4**) were used for subsequent genome-wide association study.

### Removal of outliers based on physically impossible values

Three kinds of impossible values were found in the phenotype matrix and excluded before any analysis. Growth rate, derived by curve fitting to the digital biomass time course, was observed to be negative for 14 of 227 accessions under drought and 9 of 239 under control conditions; the affected plants have projected areas and harvest weights in the normal range for the panel, so these are artifacts of the fit rather than shrinking plants. One accession (TDP_202) has Fv′/Fm′ = −2.13 and NPQ(t) = −8.17 on day 13 under control conditions, arising from an Fo′ reading larger than Fm′ (812 against 259); both are quantum or quenching quantities that cannot take these values, and the record was removed for those traits. Evapotranspiration per unit fresh weight reaches 2,017 g g⁻¹ day⁻¹ against a median of 113, in plants whose harvest fresh weight is 0.3–0.6 g. It was log-transformed and winsorized at the 98th percentile, which brings the skew from 5.2–6.6 to 0.4–0.6.

Examining evapotranspiration per unit fresh weight exposed a more basic problem with it. Its correlation with its own denominator is Spearman ρ = −0.87 under control and −0.92 under drought. In logarithms, fresh weight alone explains 83% of its variance under control conditions and 87% under drought, against 7% and 0.4% respectively for the evapotranspiration in the numerator. The quantity is therefore very largely inverse fresh weight rather than a measure of water use per unit tissue. We consequently also define a size-adjusted alternative — the residual of log evapotranspiration on log fresh weight, that is, water use beyond what plant size predicts — and report both.

All 113 numeric columns of the phenotype matrix were audited against physical bounds. Values outside them were set to missing rather than corrected: quantum yields were required to lie in [0, 1] range, and growth rate, mass, area, evapotranspiration, quenching parameters, SPAD and leaf thickness to be non-negative. Apart from the growth rates and the single fluorescence record described above, no trait used in this study contained an out-of-bounds value.

### Comparison of conditions within germplasm class

Growth rate under control and under drought was compared within each germplasm class by paired Wilcoxon signed-rank test, the accession being the unit of pairing and only accessions with a value under both conditions being used. *P*-values were adjusted across the four classes by the Holm procedure. A rank-based paired test was chosen because 1) the two conditions are not independent samples, 2) the within-accession differences depart from normality in the largest class (Shapiro-Wilk *P* = 0.004 for wild), and 3) two of the four classes contain fewer than ten accessions, where normality cannot be assessed. The same rank-based framework is used for the comparison of relative growth rate between classes (Kruskal-Wallis) and for the haplotype analysis.

#### Canopy architecture scoring

Canopy architecture was scored on the same eight phenocage experiments described above, in which every plant was imaged from seven side views on eight dates at two-day intervals. Growth habit was scored manually from a single image per plant, taken on the final imaging date of each experiment, into two categories on a single criterion: a plant whose stem was observed curling around the cage support was called **climbing**, and a plant showing no tendril development and no curling around the support was called **erect**. The scoring is categorical; the prostrate habit recognized in some *Phaseolus* classifications was not used, as the phenocage supports do not permit it to be distinguished. The bush-versus-vine contrast drawn in common bean does not apply here, because tepary accessions are indeterminate; climbing and erect therefore record whether a plant used the support, not its determinacy. A spindliness score of 0, 0.5 or 1 was recorded alongside the habit call, scoring how thin and drawn out the shoot appeared: 0 for a plant with stout stems, short internodes and dense foliage; 1 for a plant with thin, elongated stems, long internodes and sparse foliage; and 0.5 for plants between the two. The score was assigned by eye by one scorer across all experiments, so it is internally consistent but is not calibrated against a physical measurement. Of 596 scored plants, 446 were called climbing and 150 erect. Scores were obtained for 284 accessions, 273 of which carry panel phenotypes.

Because habit is partly a response to the treatment rather than a fixed property of the genotype — the control and drought plants of the same accession received the same call for 79.8% of accessions (Cohen’s κ = 0.46), and the disagreement is directional, with 35 accessions scored climbing under control and erect under drought against 18 in the opposite direction (exact McNemar *P* = 0.027)—the habit variable used throughout is the call made on the control plant alone (191 climbing, 51 erect among accessions with panel phenotypes). Results using the drought-plant call and the accession consensus are given in **Fig. S2** and do not differ.

Digital biomass is the plant pixel area (*A*) summed across the seven side views of a plant on a given date. Three further shape measures were derived from *A*, the height (*H*) and the width (*W*) of each side view and averaged over the seven views: fill fraction, aspect ratio and compactness. Fill fraction, *A*/(*H* × *W*), is the proportion of the bounding box occupied by plant pixels. Aspect ratio is *H*/*W*. Compactness is the equivalent-circle diameter divided by the bounding-box diagonal scale, 2√(*A*/π)/√(*HW*). Compactness is a monotone function of fill fraction and is reported for completeness rather than as independent evidence. Of 34,025 image rows, 592 lacked usable height or width values and returned no shape measure. These are bounding-box statistics on a silhouette: they capture how much of a rectangle a plant fills and whether it is taller than wide, and cannot resolve leaf overlap, self-shading, leaf angle or stem architecture.

### Trait contribution modeling

To identify which physiological traits contribute most to the maintenance of growth under water limitation, random forest regression models (300 trees, minimum leaf size 3) were trained on the 207 accessions with positive growth rates under both conditions. Traits with more than 40% missing values were dropped and the remainder imputed by trait median. Model performance was assessed by repeated cross-validation (5 folds × 4 repeats, 20 fits) and trait contributions by permutation importance (10 permutations per feature), in Python with scikit-learn. Two features of the design require comment, because the obvious specification does not work.

#### Target

The natural target, the ratio of growth rate under drought to growth rate under control, is heavy-tailed in this panel (skew 8.99, range 0.02–11.2; **Fig. 4A**), because accessions that barely grew under control conditions produce very large ratios. A model trained on the raw ratio does not generalize: across 20 folds its cross-validated *R*² is −0.57 ± 1.43, individual folds ranging from −5.13 to +0.71, so any single split can return an arbitrary value (**Fig. 4B**). We therefore use two stabilized targets: the rank-inverse-normal transform of the ratio, and the residual of growth rate under drought regressed on growth rate under control — that is, growth under water limitation relative to what an accession’s own well-watered vigor predicts — likewise rank-normalized. The two give nearly identical model performance (*R*Z = 0.56 ± 0.09 and 0.54 ± 0.09), and the residual is used for the models reported because it states the biological question directly and is not arithmetically coupled to its own denominator.

#### Predictors

Growth-rate components and digital biomass were excluded throughout. In the primary model we additionally excluded the harvest measurements — fresh weight, dry weight, tissue water content, and evapotranspiration expressed per unit fresh weight — because these are taken at the end of the experiment and are alternative measurements of the outcome rather than predictors of it. Their removal lowers cross-validated *R*Z from 0.54 ± 0.09 to 0.37 ± 0.17 across 72 predictors, which we take to be the honest figure. A further model restricted to traits measured on or before day 6 (32 predictors) achieves *R*² = 0.30 ± 0.16 and is genuinely predictive, in that no predictor is measured after the midpoint of the experiment. A null model with the target randomly permuted gives −0.12 ± 0.08, establishing the floor.

### Genome-wide association study (GWAS)

#### Traits and transformation

Association was tested for 104 rank-normalized measurements: digital biomass — the plant pixel area summed across the seven side views of a plant — from RGB image-based phenotyping across the treatment period (Yu *et al*., 2024), growth rate, harvest fresh weight, dry weight and tissue water content, evapotranspiration per unit fresh weight and its size-adjusted residual, daily evapotranspiration and its cumulative and median summaries, and the photosynthesis-related parameters recorded on days 6 and 13 with PhotoSynQ (Kuhlgert *et al*., 2016), each under drought and under full irrigation. Digital biomass was scanned for every imaging day from day 0, corresponding to four days before the drought pots reached their target weights, so that each locus has a pre-treatment baseline. Every trait was screened against a ladder of transformations — raw, log₁₀, square root, cube root, reciprocal, Box–Cox and Yeo–Johnson — and the one giving the highest Shapiro–Wilk statistic retained (González-Estrada and Cosmes, 2019); each trait was also transformed independently by rank-inverse-normal transformation. Only one of 112 traits is normal untransformed and only 47 reach *P* > 0.05 under the best parametric transformation, so the rank-normalized scans are reported throughout and the parametric scans are retained only as a transformation diagnostic. The selected transformations, with the distribution statistics on which the choice was based, are listed in **Tables S5–S7**. The likelihood-ratio test is reported throughout; Wald values were retained only as a diagnostic. The water-content drought/control ratio was excluded from the reported scans (likelihood-ratio inflation factor 0.36, minimum *P* = 7 × 10⁻⁶⁸).

#### Genotypes for the re-analysis

Per-chromosome VCFs were converted with plink v1.9 (Purcell *et al*., 2007), retaining SNPs with minor allele frequency above 0.05 and call rate above 0.8, giving 6,140,834 of the 6,912,813 called variants across 363 accessions. GEMMA applies the filters at run time, so the number tested varies with trait-specific missingness: 5,803,479 to 5,886,977 SNPs across the traits reported here, with 225 to 252 accessions carrying both genotype and phenotype.

#### Mixed-model association

A centered relatedness matrix was computed from the same SNPs, with missing calls mean-imputed, and each trait was tested in a univariate linear mixed model in GEMMA v0.98 (Zhou and Stephens, 2012). The scan was replicated in EMMAX (Kang *et al*., 2010) under the same filters, which returned concordant and uniformly more conservative values; those summary statistics are deposited but are not reported here.

#### Peak calling

Peaks were called as follows: every SNP reaching −log₁₀ *P* = 5, merged into one peak when lead SNPs lay within 250 kb, and peaks from different traits whose leads fell within 250 kb collapsed into one locus. For every locus reaching 6 in either scan, the strongest *other* SNP within 250 kb of the lead was recorded as a measure of linkage-disequilibrium support; loci whose lead had no neighbor above −log₁₀ *P* = 3 are treated as unsupported.

#### Linkage disequilibrium and candidate intervals

Genome-wide decay was measured on genotypes filtered to minor allele frequency ≥ 0.05 and call rate ≥ 0.8 within the 226 phenotyped accessions and thinned at random to 1.9% of SNPs, computing *r*Z for every SNP pair separated by less than 1 Mb in plink v1.9 (Purcell *et al*., 2007): 111,305 SNPs and 28.2 million pairs, mean *r*Z crossing 0.2 at 12.4 kb (**Fig. S3**). Because a single genome-wide decay distance is a poor basis for drawing intervals in a structured panel, candidate intervals were instead measured per locus on the fully analyzed genotype set. For each of the 46 suprathreshold positions, *r*Z was computed against every SNP within ±3–6 Mb, both on mean-imputed genotypes and after projecting out the first five principal components of a genomic relationship matrix built from 40,939 SNPs thinned across the genome. Each interval was taken as the contiguous block around the locus’s leads, walking outward until no SNP reached the threshold within a 100 kb gap, at *r*² ≥ 0.6 (core) and ≥ 0.3 (outer bound). Independent loci were then defined by single-linkage clustering of the 46 positions at structure-corrected *r*² ≥ 0.3, giving nine. Both uncorrected and corrected intervals are reported in **Table S12**.

#### Germplasm-class decomposition

Each lead SNP’s allele frequency was computed within the Wild, Weedy, Cultivated and Breeding germplasm classes of Table S1, and the proportion of genotype variance attributable to class taken as the between-class sum of squares over the total. Each association was then re-fitted by ordinary least squares on the full panel without covariates, with the first five principal components, with the germplasm class as a binary covariate, and separately within the Cultivated and the Wild groups alone. Call rate, minor allele frequency and heterozygosity were computed at all 46 positions as a genotype-quality check, heterozygosity being compared against a genome-wide baseline from 1,229 thinned SNPs.

#### Candidate-gene annotation and expression

Genes overlapping each structure-corrected interval were taken from the *P. acutifolius* v1.0 annotation with Arabidopsis, rice, Pfam and Gene Ontology assignments. Expression was taken from the twelve-tissue developmental atlas and leaf heat time course of Moghaddam *et al*. (2021), which use the same gene identifiers. Common bean orthologs were assigned by reciprocal best hits between the *P. acutifolius* v1.0 and *P. vulgaris* v2.1 primary-transcript proteomes (Smith–Waterman, BLOSUM62, gap open 11, extend 1), and queried against the drought experiments of Gregorio Jorge *et al*. (2020), López *et al*. (2020) and Subramani *et al*. (2023).

#### Gene-family phylogenies

All proteins carrying PF03822 (CIPK) or PF03514 (GRAS) in the *P. acutifolius* and *P. vulgaris* proteomes were collected, and the corresponding Arabidopsis families recovered from the TAIR10 representative-gene-model proteome by Smith–Waterman search against the legume family members, retaining proteins whose own best tepary hit fell inside the family (27 CIPK and 34 GRAS loci). Sequences were aligned with FAMSA (Deorowicz *et al*., 2016) and neighbor-joining trees built on p-distance over non-gap positions, with bootstrap support from 200 (CIPK) or 100 (GRAS) column-resampling replicates, midpoint-rooted.

#### Significance thresholds

A single threshold is used throughout: a Bonferroni correction over the SNPs actually analysed, which number between 5,845,456 and 5,886,977 depending on how many accessions carry the trait. At α = 0.05 this gives α_variant = 8.5 × 10⁻⁹, or −log₁₀ *P* = 8.07. Bonferroni was chosen over the alternatives because it is transparent and reproducible from the SNP count alone; it is also the more stringent of the thresholds considered, so no result reported here depends on the choice. For reference, a permutation threshold computed as in Churchill and Doerge (1994) and John *et al*. (2022, 2024a) — the phenotype vector permuted against the genotype matri× 100 times under the same model, filters and kinship matrix, with the 95th percentile of the genome-wide maxima taken — gives −log₁₀ *P* = 7.79, and effective-SNP thresholds following Li and Ji (2005) and Gao *et al*. (2008) give lower values still. Neither is used to call significance. Genomic inflation factors were estimated from a 1-in-50 genome-wide sample of SNPs.

### Data availability

All codes and computational pipelines used for re-sequencing data processing, variant calling, population genomics analyses, and GWAS were deposited on GitHub (https://github.com/Leon-Yu0320/Tepary-Genomic-Studies) and DataDryad (https://doi.org/10.5061/dryad.9w0vt4bwb). The raw sequencing reads were archived under NCBI BioProject number PRJNA1119990. The repositories holding every raw and processed dataset supporting this study are listed in **Table S9 and** https://doi.org/10.5061/dryad.9w0vt4bwb.

## Results

### A high-density SNP map for tepary bean

Tepary bean has been widely recognized for its drought tolerance, but its breeding programs require high SNP density to establish high resolution of the genetic links to be established with phenotypes recorded under water deficit. No published panel meets this high SNP density requirement, providing a rationale for this work. The tepary bean diversity panel developed here comprises 363 re-sequenced individuals: 350 accessions of the tepary diversity panel and 13 drawn from the previously published sequence data of Moghaddam *et al*. (2021). Of the 350, 162 are wild, 153 cultivated, 27 are breeding lines and 8 are weedy, so both ends of the domestication gradient are sampled in comparable numbers. The 321 accessions with collection records come overwhelmingly from Mexico and the southwestern United States, concentrated in the center of origin and in the areas where the crop is still grown (**Fig. 1A**; **Table S1**). We resequenced to 6.59–12.69× depth (**Table S2**) and recovered 6,912,813 SNPs after filtering (**Table S3**). This is a two-order-of-magnitude increase in SNP density over the panels previously reported for this species (Moghaddam *et al*., 2021; Bornowski *et al*., 2023).

**Figure 1.**
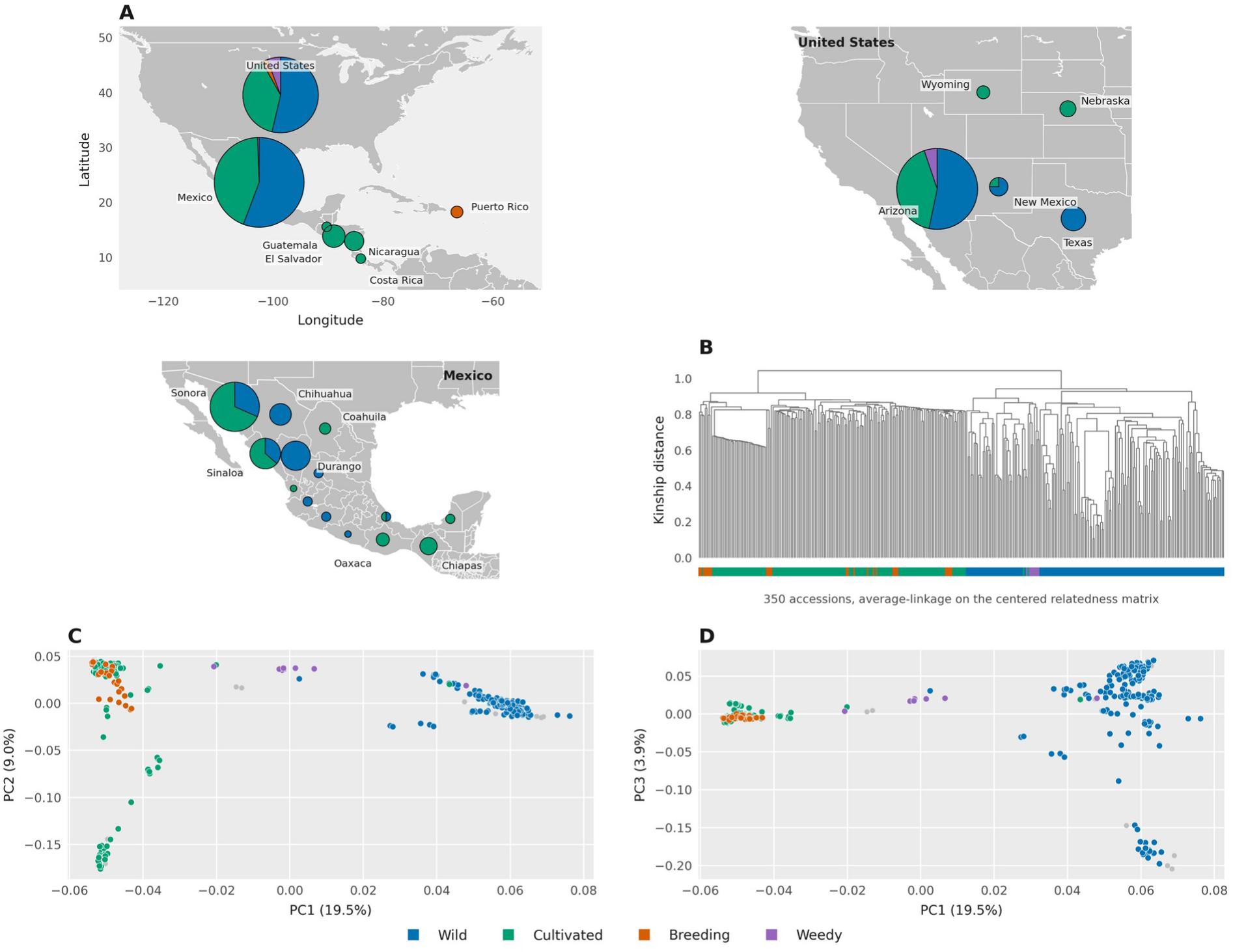
Geographic origin, relatedness and population structure of the panel. **(A)** Collection origins of the re-sequenced accessions, worldwide and at higher resolution across the United States and Mexico; pie area is proportional to the number of accessions at a location and slices give germplasm class ( **Table S1**). **(B)** Hierarchical clustering of the accessions with passport data, average linkage on the centered relatedness matrix used for association testing; the bar beneath orders accessions as in the dendrogram and colors them by germplasm class. **(C, D)** Principal component analysis of the 95,311 SNPs remaining after linkage-disequilibrium pruning (200 kb window, step 50, *r*Z < 0.2), shown as PC1 against PC2 (C) and PC1 against PC3 (D); axis labels give the variance each component explains. Point color gives germplasm class throughout.

Association studies must account for relatedness, and domestication history sets how much of the panel’s diversity is accessible within a breeding program. Hierarchical clustering by average linkage on the centered relatedness matrix separated a compact cultivated cluster from a much more internally diverse wild group (**Fig. 1B**). The breeding lines fell inside the cultivated cluster. Principal component analysis of the 95,311 SNPs remaining after linkage-disequilibrium pruning gave the same division along the first component, which explains 19.5% of the variance (**Fig. 1C, D**). ADMIXTURE gave a third, concordant view. Cross-validation error declines monotonically from *K* = 2 to *K* = 10 without an interior minimum, so no single *K* is supported. At every *K*, however, the cultivated pool resolves as one ancestry component shared with the breeding lines, and wild accessions divide among the remainder (**Fig. S1**). Wild germplasm therefore retains substantially more diversity than the cultivated pool. This is consistent with a single domestication event (Garvin and Weeden, 1994) and with results reported for both previous panels.

### Domestication increased vigor without eroding drought resilience

To evaluate whether domestication results in reduced drought resilience in tepary bean, we compared growth across germplasm classes of plants grown under control and drought stress treatments, corresponding to 60 and 10% of soil water holding capacity (SWHC) respectively. Drought reduced growth rate in every class (**Fig. 2A-C**). Cultivated and breeding lines grew roughly twice as fast as wild accessions under both regimes (**Fig. 2C**). The proportional reduction, however, did not differ between the four classes (Kruskal–Wallis *P* = 0.29; **Fig. 2D**). Similar trends were observed for other size-related parameters, such as fresh weight at harvest and tissue water content (**Fig. S4**). Fresh weight at the terminal point of the experiment confirmed that every class lost weight under drought (paired Wilcoxon, Holm-adjusted *q* = 0.031 to 1.4 × 10⁻¹⁸), and under full irrigation the class ranking followed the growth rates, from breeding lines down through cultivated and weedy to wild (**Fig. 2E**). Although the comparison between the classes remains unbalanced, these results suggest that breeding selection for vigor did not come at the cost of drought resilience.

**Figure 2.**
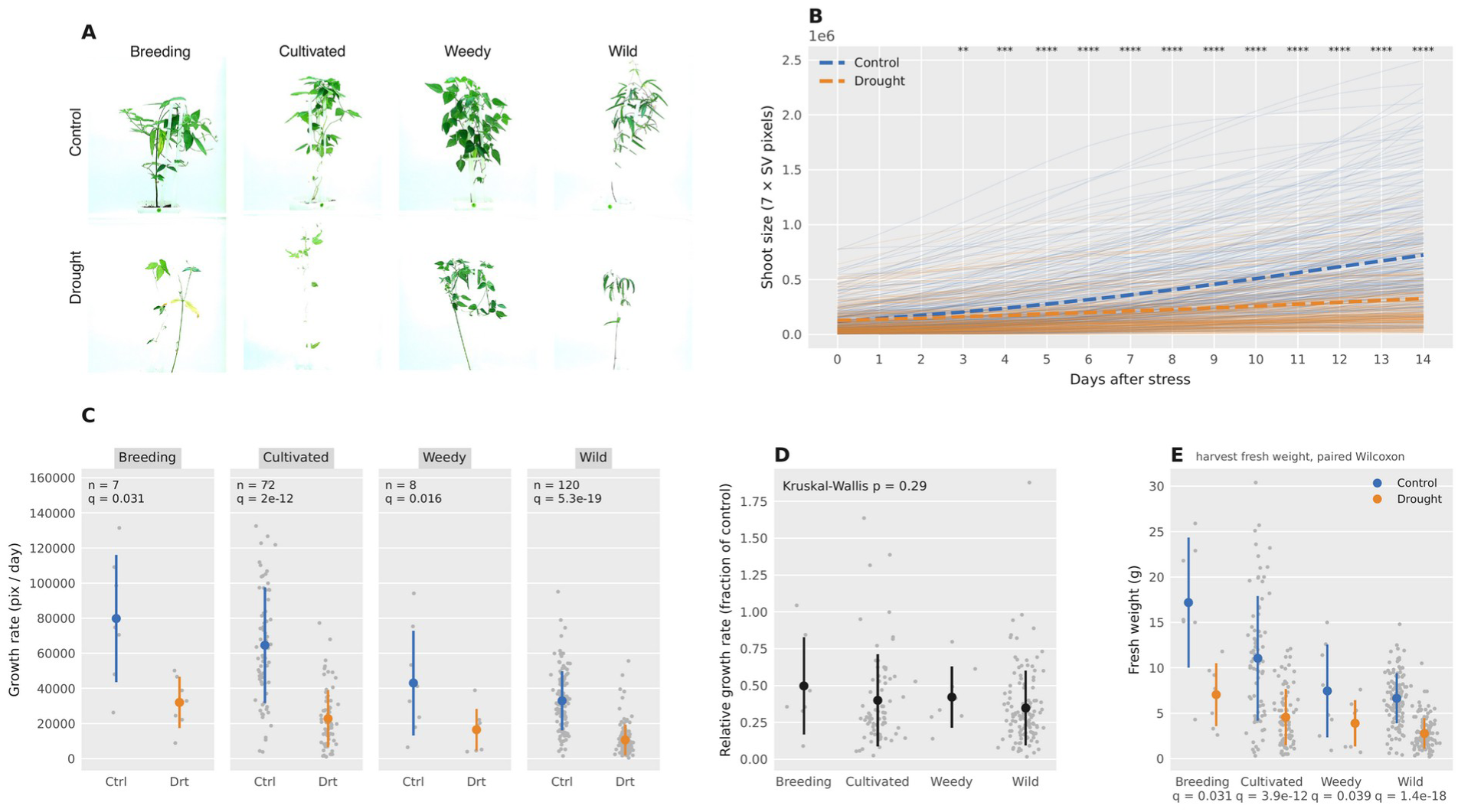
Growth and harvest traits under control and drought, by germplasm class. **(A)** Representative plants under control (60% soil water holding capacity, SWHC, top) and drought (10% SWHC, bottom). **(B)** Shoot size (sum of seven side-view projections, pixels) against days after stress imposition for every plant; thin lines, individual plants colored by treatment; thick dashed lines, treatment means. Asterisks mark the significance of the treatment difference on each day (Mann–Whitney). **(C)** Growth rate (pixels day⁻¹) under control and drought within each germplasm class; gray points, individual accessions; colored point and bar, mean ± standard deviation (SD). *q*, paired Wilcoxon signed-rank test within class, Holm-adjusted across the four classes; *n*, accessions with a value under both conditions. **(D)** Relative growth rate (drought/control ratio) by germplasm class, with the Kruskal–Wallis test of a between-class difference. **(E)** Harvest fresh weight under control and drought within each germplasm class; conventions and test as in (C).

### Evapotranspiration is largely a function of plant size

Evapotranspiration often explains plant performance under water stress (Morton *et al*., 2024), and the AAWESMO gravimetric platform both holds the deficit at a reproducible level across genotypes and records daily water loss. The collected dataset allowed us to explore whether daily evapotranspiration is a property of the genotype or of plant size. Evapotranspiration, measured as daily water loss over the 14-day treatment, separated the two regimes within a day (**Fig. 3A**). The same signal is visible using the direct leaf measurements with PhotoSynQ: accessions with cooler leaves on day 13 used more water over the course of experiment (**Fig. 3B**). Drought reduced evapotranspiration within every germplasm class, but the proportional reduction did not differ between classes (**Fig. 3C, D**). Growth rate showed the same pattern. The classes differ several-fold in size, and evapotranspiration per unit fresh weight ran exactly opposite to that gradient. It was highest in wild accessions and lowest in breeding lines (**Table S4**). That inverse gradient is not water-use efficiency *per se*. We regressed log evapotranspiration on log fresh weight. Fresh weight explained the great majority of the variance in observed evapotranspiration per fresh weight, and evapotranspiration itself explained almost none. The index is therefore very largely the reciprocal of plant size. We analyzed the residual instead, which is the water lost beyond what size predicts (**Fig. 3E**). On that scale the class differences did not vanish, but they changed character. Breeding lines used about 30% less water than plants of their size were predicted to, while wild accessions used what their size predicted. The classes remained distinguishable.

**Figure 3.**
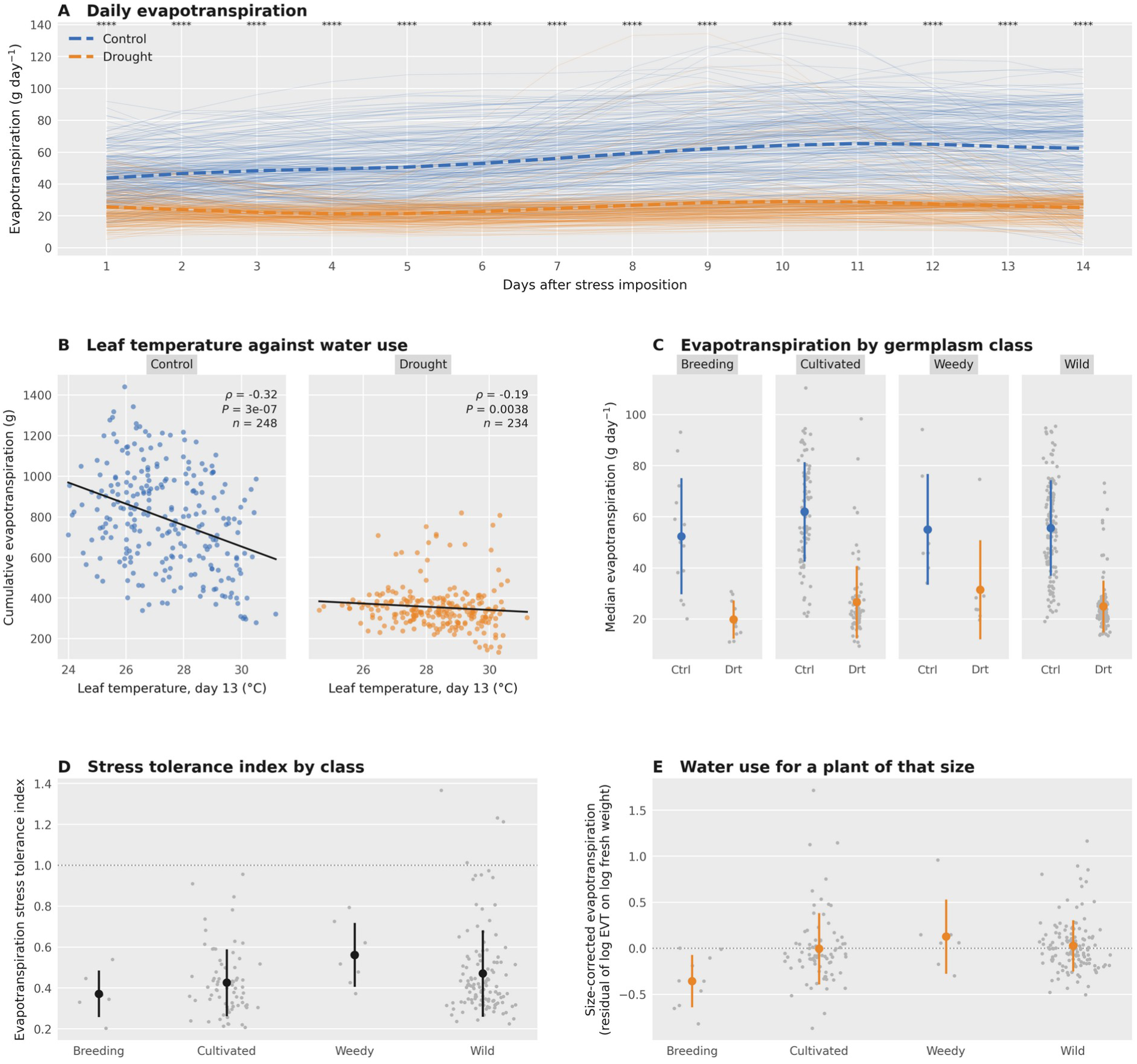
Evapotranspiration, its leaf-level correlate, and what remains of it after plant size is removed. **(A)** Daily evapotranspiration (g day⁻¹) for every plant across the 14 days of treatment; conventions as in Fig. 2B. **(B)** Cumulative evapotranspiration against leaf temperature measured on day 13, under control (left) and drought (right); line, least-squares fit; ρ, Spearman rank correlation, with its *P* and the number of accessions. **(C)** Median evapotranspiration under control and drought within each germplasm class; gray points, individual accessions; colored point and bar, mean ± SD. **(D)** Evapotranspiration stress tolerance index (median drought / median control) by germplasm class; dotted line, no reduction. Values above the 99th percentile were excluded as uninterpretable. **(E)** Size-corrected evapotranspiration under drought — the residual of log median evapotranspiration regressed on log harvest fresh weight — by germplasm class; dotted line, the water use predicted for a plant of that size. Positive values are accessions using more water than their size predicts.

#### Water use predicts the maintenance of growth

Canopy architecture is the only trait class that can be read from a single image, which would make it the cheapest selection criterion available to a breeding program. We therefore asked whether it carries information about drought performance. We scored it two ways on the same phenocage series. First, we scored the growth habit manually as climbing or erect from one image per plant. Second, we automatically extracted four measures from the silhouette: digital biomass, the plant pixel area summed across the seven side views, and three shape measures averaged across those views — fill fraction, aspect ratio and compactness (**Table S10**). We tested each against water use and against the drought response itself (**Fig. 4A**). Larger, fuller canopies used more water in absolute terms. Once the differences in fresh weight were corrected for, the pattern reversed: for a given plant mass, a more compact canopy used less water (**Table S11**). We define the size-independent maintenance of growth as the residual of growth rate under drought regressed on growth rate under control, so that it measures how much better or worse an accession grew under water deficit than its own control-condition vigor predicts. Against that measure, both canopy measures and growth habit scored weakly (**Fig. S2**). Climbing and erect accessions did not differ in the proportional maintenance of growth, and habit was largely a proxy for wild-versus-cultivated status. Canopy architecture therefore predicted how much water a plant uses, but not their relative drought performance.

**Figure 4.**
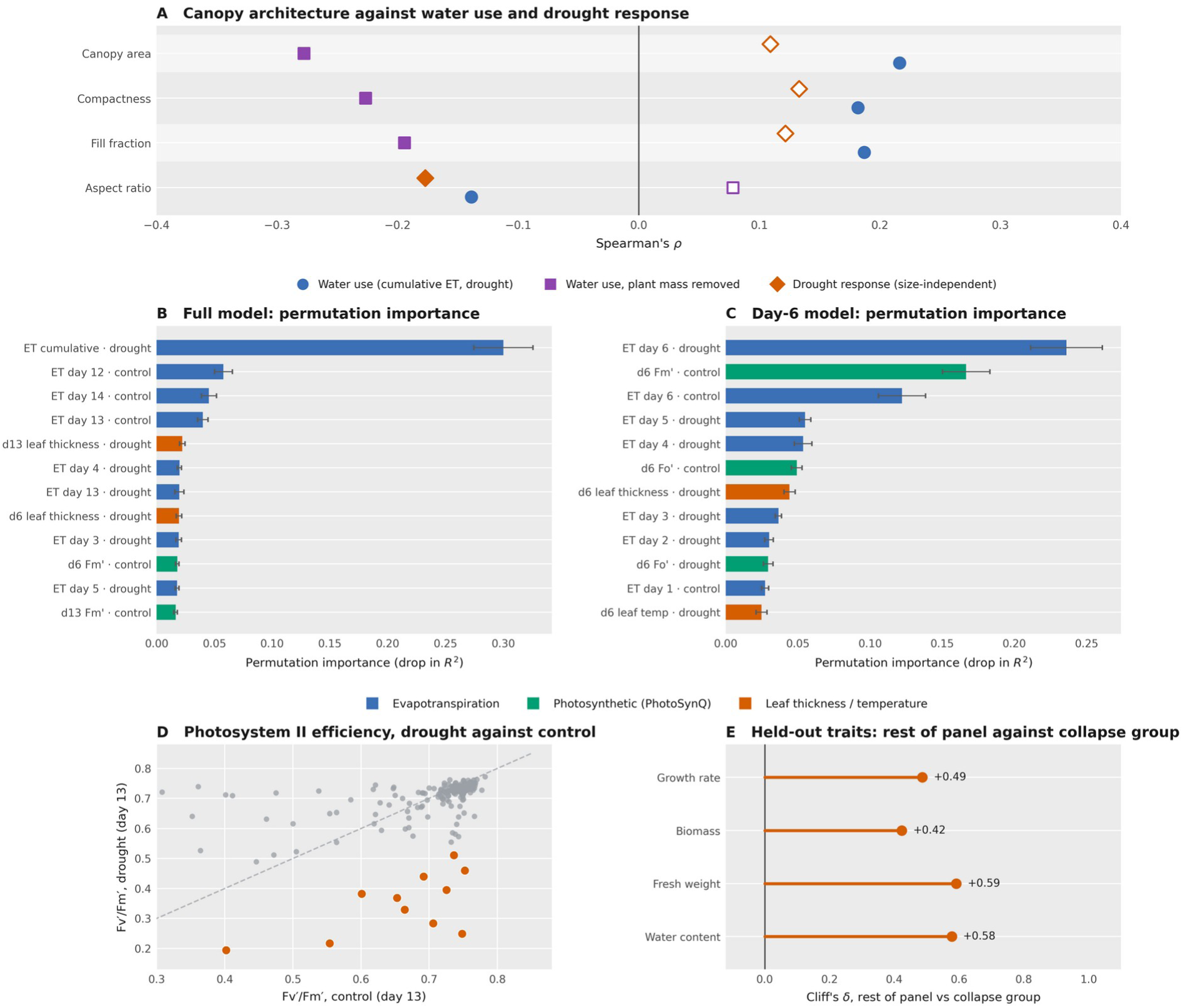
Canopy, water-use and photosynthetic traits against the maintenance of growth under drought. **(A)** Each canopy measure against three targets on a common rank-correlation axis: cumulative evapotranspiration under drought, the same with harvest fresh weight partialed out, and the size-independent drought response (the residual of drought growth rate regressed on control growth rate). Filled SNPs, *P* or *q* < 0.05; *n* = 209–238. **(B)** Permutation importance of the twelve strongest predictors in the random forest trained on the size-independent drought response with harvest measurements excluded (72 predictors); bars colored by trait family, error bars SD over ten permutations. **(C)** The same for the model restricted to the 32 traits measured on or before day 6, in which no predictor postdates the outcome. **(D)** Day-13 Fv′/Fm′ under drought against the same accession’s control value; orange, the twelve accessions separated by *k*-means clustering on the photosynthetic drought response alone (silhouette 0.75); dashed line, parity. One accession with an out-of-range stored value is omitted. **(E)** Cliff’s δ between the rest of the panel and those twelve accessions, for four traits held out of the clustering, each expressed as the residual of the drought value on the control value. Positive δ means the rest of the panel retained more. The growth comparison uses the six accessions of the group with a positive growth rate under both treatments.

Traits measured at single leaf level with PhotoSynQ behaved differently on the two measurement days, although none separated the germplasm classes (**Fig. S5**). Warmer leaves used more water per unit plant mass early in the experiment, but that relationship was not observed anymore by day 13. Leaf thickness showed the opposite pattern. It had no early relationship, but turned strongly negative under drought by day 13. For a given plant mass, thicker-leaved accessions were therefore the more conservative water users during longer exposure to drought. As correlations test one trait at a time, we subsequently trained a random forest on all traits jointly against the size-independent maintenance of growth. The model predicted the outcome well above chance. Cumulative evapotranspiration under drought dominated it, with a permutation importance five times that of the next-ranked variable (**Fig. 4B**). However, we also wanted to explore the predictive potential of traits scored at earlier times, as the predictors measured at or near harvesting time cannot support a selection decision made during the experiment. We therefore refit the model using only traits recorded on or before day 6, and the model retained genuine predictive power (**Fig. 4C**). Although drought-to-control ratio is the intuitive target, but it is so heavy-tailed in this panel that a model trained on it did not generalize (**Fig. S6**). Therefore, we used the size-independent residual throughout.

Every analysis above assumes that the panel responds along common axes and differs only in degree. We tested that assumption by letting accessions sort themselves on one block of traits and then testing them on another. Clustering on the photosynthetic drought response alone separated twelve accessions from the rest. In those twelve, photosystem II efficiency halved under drought and non-photochemical quenching rose by an order of magnitude (**Fig. 4D**; **Fig. S7**). The grouping used no growth, size or water data, yet predicted all of it (**Fig. 4E**), and it is independent of germplasm class: the twelve comprise seven wild, three cultivated and one breeding accession, with one not recorded, in proportions close to the panel as a whole (**Table S16**). Twelve accessions in a single experiment make this an observation that requires replication. The above results reveal two patterns of drought sensitivity within tepary bean diversity panel: an accession can arrive at poor growth maintenance either by using water freely and running out, or by shutting photosynthesis down early, and a panel-wide correlation cannot distinguish the two.

### Association mapping resolves nine loci, and drought and control barely overlap

Having established which traits correspond the maintenance of growth (**Fig. 4**), we further explored which regions of the genome correspond to observed variation in recorded phenotypes. We tested association for 104 rank-normalized measurements (**Tables S5–S7**) across 5.8 million SNPs in a linear mixed model, using the likelihood-ratio test throughout. Each trait used for GWAS with data from 216 to 252 accessions, depending on how many had a value for it ( **Table S7**). Genomic inflation lay between 0.92 and 1.07 (**Fig. S8**), suggesting that neither residual population structure nor over-correction by kinship distorted the GWAS test statistics. Against a Bonferroni threshold over the SNPs analyzed (−log₁₀ *P* = 8.07; Methods), 113 SNPs at 46 distinct positions exceeded significance threshold. Since significant SNPs positioned within LD cannot be considered as independent, we further consolidated 46 positions by LD measured on the analyzed genotypes and corrected for population structure. Joining SNP positions with *r*² ≥ 0.3 resolved our significantly associated SNPs into nine independent loci (**Fig. 5A**; **Table S12**). The nine loci are what this panel had the power to detect, and the sub-threshold peaks — 3,340 above−log₁₀ *P* = 5 and 595 above 6 — may still hold regions contributing to these traits (**Fig. 5A**; **Supplementary Results S1**). The loci are not distributed in proportion to SNP density, so the absence of association elsewhere is not an absence of SNPs (**Fig. S3**). **Figure S9** provides more detailed overview of which traits nominate which region.

**Figure 5.**
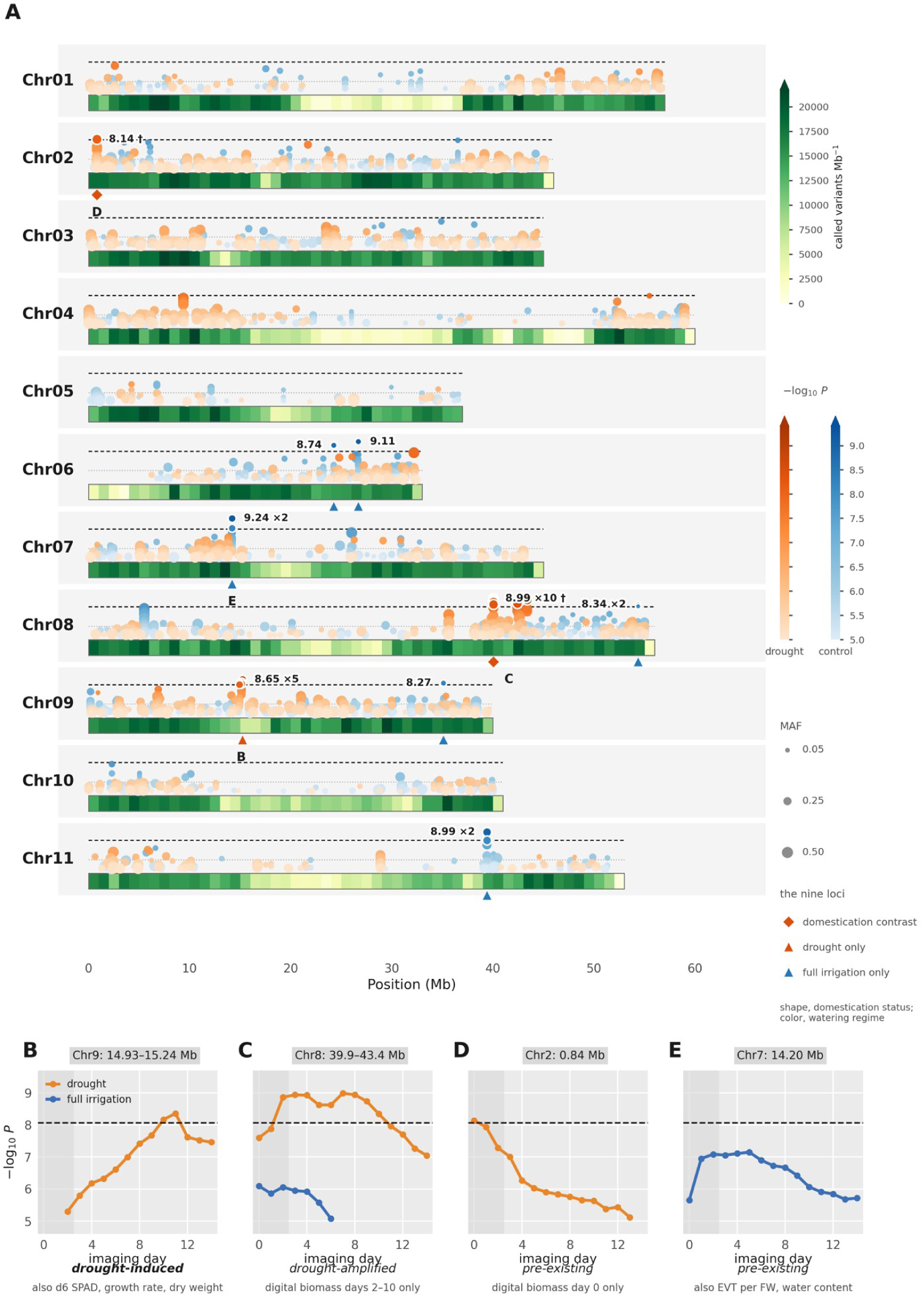
Every association in the genome, and when each locus appears. **(A)** Each chromosome carries a SNP-density ideogram (green, called variants per megabase) with a Manhattan strip above it. Every independent peak reaching −log₁₀ *P* = 5 in any of the twenty-one rank-normalized measurements is one mark; peaks whose lead SNPs lie within 250 kb are treated as one locus. Within a strip, height and color both give −log₁₀ *P* (5 at the bottom of the strip, 9.4 at the top), orange is used for traits measured under drought and blue for traits measured under full irrigation, and marker area gives the lead SNP’s minor allele frequency. Dashed line, the Bonferroni threshold (−log₁₀ *P* = 8.07; Methods); dotted line, −log₁₀ *P* = 6. Peaks clearing the threshold are outlined in white and labeled with the best *P* at that locus and, where more than one measurement peaks there, the number of measurements. Labels are grouped by the nine linkage-disequilibrium-resolved loci. Beneath each chromosome, one SNP per locus gives its category (**Table S12**): a diamond marks a domestication contrast and a triangle a trait-specific locus, and the color gives the regime the locus was nominated under, orange for drought and blue for full irrigation. A dagger on a label marks the same two domestication loci. Letters mark the four loci shown in panels B–E. **(B–E)** Onset of four representative loci. Digital biomass was imaged from day 0, four days before the drought pots reached their target weights, so the association can be run for each imaging day separately and every locus has a pre-treatment baseline; the grey band marks the pre-treatment days. Each point is the best−log₁₀ *P* within the region for digital biomass on that day, orange under drought and blue under full irrigation; dashed line, the Bonferroni threshold. Beneath each panel is the onset class and the other measurements that reach the threshold in that region. **(B)** Chromosome 9 at 14.93–15.24 Mb is undetectable before the treatment and rises monotonically through it: drought-induced. **(C)** The chromosome 8 block is already elevated on day 0 but holds above the threshold under drought while decaying to below detection under full irrigation: drought-amplified. **(D, E)** Chromosome 2 at 0.84 Mb and the control-side chromosome 7 region at 14.20 Mb peak at or near day 0 and decay thereafter: pre-existing differences in seedling size.

The LD correction mattered most for loci on chromosome 8. As called, five of its regions fell inside a single block of 7 Mb at *r*² ≥ 0.6, and the chromosome 2 region spanned 1.6 Mb. Once population structure was removed, those blocks collapsed to between 0 and 182 kb, so the long-range component was related to ancestry rather than linkage. The remaining intervals, which did not collapse with ancestry, ranged from 2.5 kb at Chr9:14.23 Mb and 26 kb at Chr11:39.42 Mb to 518 kb at Chr9:14.93–15.24 Mb. One region, Chr8:54.36 Mb, has no SNP reaching *r*² ≥ 0.2 anywhere within 3 Mb. Its association rests on a single uncorroborated SNP, and we treat it as provisional. Two caveats attach to these intervals. First, removing principal components also removes real disequilibrium wherever a causal haplotype is itself correlated with structure. The true interval for each locus is therefore bracketed by the uncorrected and corrected estimates, and we report both. Second, *r*Z is attenuated by genotyping error. The mean between SNPs less than 1 kb apart is 0.22 genome-wide, so these intervals err toward being too wide rather than too narrow.

Since this panel spans wild, weedy, cultivated and breeding material, and plant size and growth differ systematically among those classes (**Fig. 2**), an association with a growth trait may report domestication rather than a trait-specific allele. We partitioned each lead SNP by germplasm class, and re-tested each association within the cultivated and the wild group separately (**Table S12**). For the chromosome 8 block, the derived allele was at a frequency of 0.91–1.00 in wild accessions and 0.06–0.09 in cultivated ones, and germplasm class accounted for 69–86% of the variance in genotype. For Chr2:0.84 Mb the corresponding figures were 0.95, 0.02 and 85%. Neither survived within either group (−log₁₀ *P* ≤ 2.9). These two loci therefore report domestication rather than a drought-specific response. On the other hand, for the two chromosome 11 leads class accounted for only 2.7% and 4.5% of the genotype variance, and they were the only associations recovered separately in cultivated accessions (−log₁₀ *P* 3.9 and 4.9) and in wild accessions (6.1 and 5.1). Six remaining loci lay between these two extremes. Their minor allele was confined to cultivated germplasm, and they replicated within that group alone. Chr9:14.23 Mb led them, reaching −log₁₀ *P* = 6.4 in 65 cultivated plants and 9.3 in the full panel after PC correction. It was the least attenuated association in the study. Genotype quality does not explain this structure: heterozygosity at the 46 positions averaged 0.003 against a genome-wide 0.009 and call rates were 0.81–0.98, the lowest four being the chromosome 11 leads (**Table S12**).

Since we imaged plants from the moment that the plants were exposed to altered watering regime, and it took on average four days before the drought pots reached their target weights, we could therefore run the association separately for each imaging day, which gave every locus a pre-treatment baseline. This separated three kinds of loci (**Fig. 5B–E**). The chromosome 9 region was undetectable on days 0 and 1. It reached −log₁₀ *P* = 5.3 on day 2 and then climbed to 8.36 by day 11. We call this class **drought-induced**. The chromosome 8 block stood at 7.60 on day 0 and cleared the threshold by day 2. It then held between 8.3 and 9.0 for the rest of the treatment, while the same region under full irrigation decayed from 6.09 to below detection by day 7. Water limitation therefore sustained and amplified a difference that was already present, and we call this class **drought-amplified** rather than drought-induced. The **pre-existing** associations are illustrated by locus on chromosome 2 region at 0.84 Mb, which reached its maximum of 8.14 on day 0, before any pot weight was able to reach drought conditions, and decayed monotonically to 5.1 by day 13. The control associations on chromosome 7 region at 14.20 Mb belongs to it as well, peaking on days 1 to 5 and decaying thereafter.

Across the nine loci, three were already present before the treatment began, five appeared after it, and one had no digital biomass series. Three were significant only under drought and six only under full irrigation. None was significant under both (**Fig. 5A**; **Table S12**). Where a region was detectable in both regimes at all, the gap was large: about three log units for the chromosome 8 block, and 3.45 in the opposite direction on chromosome 7. The separation is therefore not an artifact of where the threshold falls. These results indicate that the genetic basis of growth under water limitation is largely separate from that of growth when water is not limiting. Read together with the germplasm-class analysis above, the chromosome 2 and chromosome 8 patterns are those of alleles that distinguish wild from cultivated plants and that separate the classes most strongly under stress, rather than of drought-response alleles as such.

### Two loci report domestication rather than a drought response

The larger of the two is a block on chromosome 8 between 39.89 and 43.39 Mb. Its best SNP reached 8.99 at Chr8:40,059,877, at a minor allele frequency of 0.48 and an effect size of *β* = −0.58, and nine measurements reached it. Association was elevated continuously across the whole block. The lead SNPs sat at a near-constant allele frequency of 0.48 ± 0.03 with a consistent effect sign, and both frequency and signal fell away abruptly beyond 44.5 Mb. This suggests one long haplotype rather than of several independent loci (**Fig. S3**). Since we already discussed this locus to track germplasm class (**Fig. 5A**; **Table S12**), it admits no gene nomination.

The other locus that tracks domestication lies at the far end of chromosome 2. Its single significant SNP, Chr2:842,944, reached −log₁₀ *P* = 8.14 for digital biomass on day 0 under drought, at a minor allele frequency of 0.45 and a call rate of 0.93. The derived allele was at a frequency of 0.95 in wild accessions and 0.02 in cultivated ones, germplasm class accounted for 85% of the variance in genotype, and the association survived in neither class alone. Like the chromosome 8 block, it was already present on the first day of imaging, before the drought pots reached their target weights, so it describes a difference in seedling size between wild and cultivated plants rather than a response to water deficit.

### One locus is specific to water limitation

Of the three loci nominated under drought and by no trait measured under full irrigation, two are the domestication contrasts above. The third is specific to water limitation in the stricter sense, and the three together carry 98 of the 113 significant SNPs (**Fig. 5A**; **Table S12**). The strongest interpretable locus lies on chromosome 9 between 14.93 and 15.24 Mb, with its highest SNP at Chr9:15,242,351 (−log₁₀ *P* = 8.65, minor allele frequency 0.28, *β* = +0.58). Under drought this locus was associated with five measurements — chlorophyll index on day 6, growth rate, dry weight, and digital biomass on days 10 and 11 — and with none under full irrigation (**Fig. 6A, B**). Because the chlorophyll index was measured on a separate instrument and is independent of plant size, it independently confirms the association.

**Figure 6.**
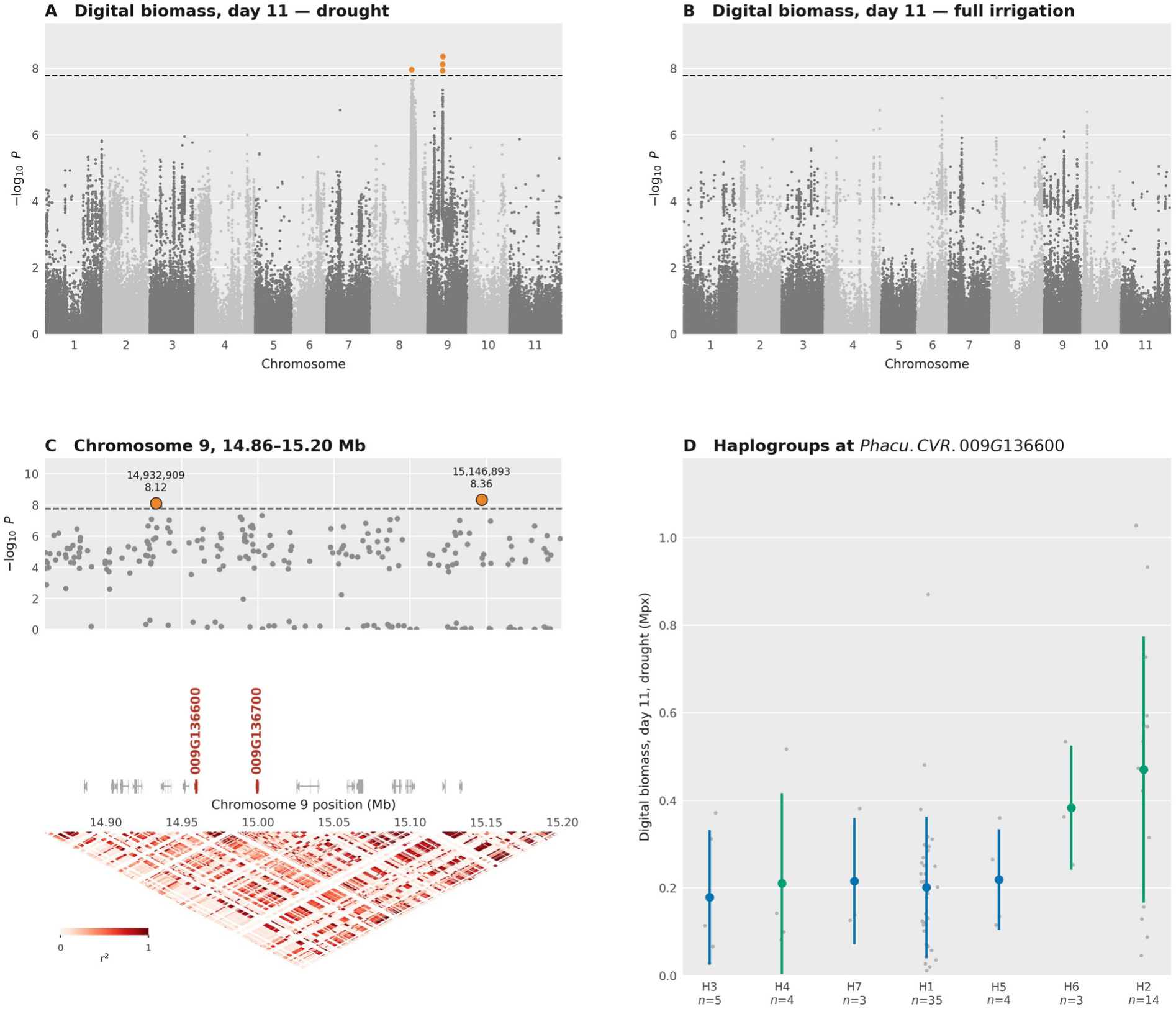
Digital biomass under drought and full irrigation, and the chromosome 9 locus. **(A, B)** Manhattan plots for digital biomass on day 11 under drought (A) and full irrigation (B), from a linear mixed model in GEMMA over 5.80–5.89 M SNPs, likelihood-ratio test, rank-normalized traits. Dashed line, the Bonferroni threshold (−log₁₀ *P* = 8.07; Methods); colored points exceed it. **(C)** The chromosome 9 locus, 14.86–15.20 Mb. *Upper*, association for the trait in (A); orange, the two lead SNPs, labeled with position and −log₁₀ *P*; dashed line, the permutation threshold. *Middle*, gene models for every annotated gene in the window (primary transcript; thick boxes, coding exons; thin boxes, untranslated regions; arrowhead, strand), with the two candidate copper radical oxidases (*Phacu.CVR.009G136600* and *Phacu.CVR.009G136700*) labeled and colored and all other genes drawn in gray. *Lower*, pairwise linkage disequilibrium (*r*Z) among SNPs across the same window, drawn so that each SNP sits beneath its position on the map. **(D)** Haplogroups defined by the genotype combination across *Phacu.CVR.009G136600* plus 3 kb of flanking sequence (24 SNPs; haplogroups of *n* ≥ 3 retained), against digital biomass on day 11 under drought. Gray points, individual accessions; colored point and bar, mean ± SD, colored by the haplogroup’s majority germplasm class; *n* beneath each haplogroup.

### Six loci are specific to full irrigation, and four of them report plant size

Four of the six loci nominated under full irrigation were led by evapotranspiration per unit fresh weight, a trait that is very largely the reciprocal of plant size (**Fig. 3E**): Chr6:24.27 Mb, Chr6:26.68 Mb, Chr7:14.20 Mb and Chr9:35.10 Mb. The size-adjusted scan returned no suprathreshold SNP anywhere in the genome, so none of the four survived it. We therefore report these four as size loci rather than as water-use loci.

The remaining two loci nominated under full irrigation were Chr8:54.36 Mb and the chromosome 11 locus at 39.42–40.02 Mb. The chromosome 11 locus is the one association in this study that is almost free of population structure, and it is treated with its candidate genes below. Chr8:54.36 Mb has no SNP reaching *r*² ≥ 0.2 anywhere within 3 Mb, so its association rests on a single uncorroborated SNP and we treat it as provisional.

### Three loci carry candidates whose expression and published mechanism match the trait

A genome-wide scan delivers an interval, not a SNP. We therefore annotated every gene in every structure-corrected interval (322 genes; **Table S14**). We subsequently explored which of these genes were expressed in the tissue and condition in which we measured the trait. We took expression from a twelve-tissue tepary developmental atlas and a leaf heat time course, both of which carry the same gene identifiers (Moghaddam *et al*., 2021). We also assigned common bean orthologs by reciprocal best hits and queried three common bean drought experiments (Gregorio Jorge *et al*., 2020; López *et al*., 2020; Subramani *et al*., 2023). Three genes satisfied position, interval and expression together (**Table S15**).

*Phacu.CVR.009G130300* encodes a cytokinin oxidase/dehydrogenase. It overlaps the lead for chlorophyll index on day 6 under drought, inside a structure-corrected interval of 2.5 kb, and its association was the least attenuated in the panel when we removed structure. The gene was near-silent in well-watered tepary tissue. Its common bean ortholog, however, was expressed and leaf-biased in droughted field material. The gene is therefore expressed in the tissue and condition in which we measured the trait, and not in unstressed chamber plants. Cytokinin oxidase catalyzes the irreversible degradation of cytokinin. Lowering its activity in the shoot raises cytokinin and delays drought-induced senescence, retaining both chlorophyll and photosystem II efficiency. This has been shown by partial silencing in sugarcane (Masoabi *et al*., 2023), by inducible overexpression in Arabidopsis (Přerostová *et al*., 2018), and in apple, where *MdCKX5* acts as a negative regulator of drought tolerance and its silencing improves tolerance (Jia *et al*., 2025). These results suggest that variation at this gene acts on chlorophyll retention under drought.

*Phacu.CVR.009G274300* encodes a CBL-interacting protein kinase and overlaps tfte lead for evapotranspiration per unit fresft weigftt at Cftr9:35.10 Mb. Its ortftolog sat in tfte 97tft–98tft percentile of tfte drougftted common bean transcriptome. It was also down-regulated by drougftt in aerial tissue (log₂ fold cftange −2.88, *P* = 0.0012). The CBL1/9–CIPK1 module is a convergence point for abscisic-acid-dependent and -independent osmotic stress signaling (D’Angelo *et al*., 2006). It regulates drought negatively, by phosphorylating PYL abscisic-acid receptors and suppressing their activity, and *cipk1* mutants are drought-resilient (Zhang *et al*., 2023). *Phacu.CVR.007G126400* encodes the photosystem II antenna subunit LHCB5/CP26. It lies 8.3 kb from a lead for the same trait and for water content at Chr7:14.20 Mb. Principal-component correction left that interval unchanged, so it reflects physical linkage rather than admixture. The gene was the most abundant transcript among the candidates (1,155 FPKM in leaf) and was induced by drought in common bean leaf (log₂ fold change +1.66, *P*adj = 2.0 × 10⁻⁷). Down-regulating any of the six Arabidopsis *LHCB* genes, *LHCB5* included, blunts the stomatal response to abscisic acid and reduces drought tolerance (Xu *et al*., 2012). The interval holds twenty other genes, so this nomination rests on expression and pathway more than on position.

Best-hit annotation within a large gene family names a relative rather than a clade. We therefore placed the kinase and transcription-factor candidates in family phylogenies. We aligned all CIPK-domain (PF03822) and GRAS-domain (PF03514) proteins of tepary bean, common bean and Arabidopsis, and built neighbor-joining trees with bootstrap resampling (**Fig. S12**). *Phacu.CVR.009G274300* pairs with *Phvul.009G220200* at 100% support within a four-member clade, also at 100% support, completed by *Phacu.CVR.003G163400* and *Phvul.003G121600* — the quartet expected of a single ancestral gene after the legume whole-genome duplication. That quartet is sister, again at 100% support, to a clade containing exactly two of the twenty-seven Arabidopsis CIPKs: **CIPK1** (AT3G17510) and AT1G48260. The GRAS candidate *Phacu.CVR.011G176400* likewise pairs with *Phvul.011G142300* at 100% support inside a legume quartet, and that quartet falls within an eighteen-member clade at 100% support whose only Arabidopsis members are PAT1 (AT5G48150), SCL5 (AT1G50600), SCL13 (AT4G17230) and SCL21 (AT2G04890) — that is, the PAT1 subfamily of GRAS.

The locus with the strongest genetics yielded the weakest mechanism. Chr11:39.42–40.02 Mb was the only locus segregating in both wild and cultivated germplasm and recovered separately in each. Its intervals of 26 and 71 kb are the tightest we obtained. Two genes lie within them. *Phacu.CVR.011G176300* encodes a receptor kinase whose 5′ end is 1.75 kb from the lead. *Phacu.CVR.011G176400* encodes a GRAS-family transcription factor 22 kb away, which the phylogeny places inside the PAT1 subfamily at 100% support. Both were expressed, and the second was root-enriched in droughted common bean. Neither has a published function in water relations.

Expression evidence was lacking for four candidates that position alone would have favored. These were a tandem pair of nuclear factor Y subunits 6 kb from a chromosome 6 lead, a MYB-family factor overlapping a second chromosome 6 lead, an array of five stress-upregulated Nod19 genes beside the chromosome 11 locus, and an amidohydrolase 264 bp from the second chromosome 11 lead. None was detected across the twelve tepary tissues or the heat time course. The first two were absent from the droughted common bean data, or in its first percentile (**Table S15**). We report them because each would otherwise be an attractive nomination. Because the atlas covers a single genotype in tissues that do not include water deficit, this is an absence of supporting evidence rather than evidence of absence, and these genes are not excluded.

### The chromosome 9 locus carries a tandem pair of copper radical oxidases

At one of the nine loci the candidate genes point at a specific biochemistry: the handling of reactive carbonyl compounds, of which methylglyoxal is the best characterized in plants. We followed that locus further. The assignment is positional, and the genes are candidates rather than identifications. **Figure S10** gives full gene models and SNP consequences.

At 309 kb, with a structure-corrected core of 518 kb, the region is narrow enough to nominate genes at all (**Fig. 6C**). It is not, however, a separate locus from the cytokinin oxidase described above (**Table S15**): the two lead SNPs are correlated at *r*Z = 0.40 after structure correction and cluster together, so the oxidase pair is the second candidate at one chromosome 9 locus rather than the only candidate at its own. Two lead positions define the region. Chr9:15,146,893 led for digital biomass and fresh weight, and Chr9:14,932,909 for dry weight. Water content also reached the region, but no trait measured under full irrigation did. Twenty-one annotated genes lie within 150 kb (**Table S8**). The candidates of interest are *Phacu.CVR.009G136600* and *Phacu.CVR.009G136700*. They form a tandem pair, carry the copper-radical-oxidase domains PF07250 and PF09118, and are annotated as glyoxal-oxidase-related. Haplogroups defined across *Phacu.CVR.009G136600* separated on digital biomass under drought (**Fig. 6D**), but that test was sensitive to how we drew the window. Three further observations weigh against the pair. The only gene in the region that contains a significant SNP, rather than merely neighboring one, is the B-box zinc finger *STH2*. Of the two oxidases, only *Phacu.CVR.009G136700* carries a significant SNP, and only it was expressed in droughted common bean root. *Phacu.CVR.009G136600*, on which the haplogroup analysis in **Fig. 6D** is built, was silent across all twelve tepary tissues and fell in the bottom tenth of the droughted common bean distribution (**Table S15**). *STH2* was the most strongly root-partitioned gene among all candidates we examined under drought. Neither oxidase is therefore the best-supported candidate at this locus once expression is taken into account.

## Discussion

Tepary bean is cited as a model of crop drought tolerance more often than it has been measured as one. Its resilience is well attested agronomically across arid North America, but the phenotypic record has consisted largely of endpoint performance under uncontrolled field stress (Mhlaba *et al*., 2018; Mwale *et al*., 2020). Here we supplied both halves of what a quantitative genetic analysis requires: a SNP map two orders of magnitude denser than the panels previously reported for this species, and a controlled record of growth, evapotranspiration and photosynthesis across 282 accessions held gravimetrically at a constant water deficit.

Cultivated and breeding lines grew roughly twice as fast as wild accessions without losing proportional drought resilience. This is not what the domestication literature leads one to expect. Bottlenecks are held to erode the standing variation on which adaptation acts (Warschefsky *et al*., 2014; von Wettberg *et al*., 2018). In *Phaseolus* specifically, wild common bean holds drought-adaptive allelic variation structured by environment of origin that cultivated germplasm largely does not (Cortés and Blair, 2018). In chickpea, another selfing legume, domestication left cultivated material with a measurably narrower transcriptional drought response than its wild progenitor (Moenga *et al*., 2020). Tepary appears to have lost the diversity without losing the drought-resilient phenotype, and to have taken a different route from common bean, where most domestication traits were fixed with little loss of genome-wide heterozygosity and the marked erosion within cultivars arrived only with intensive recent selection (Trucchi *et al*., 2021). Erosion of diversity therefore need not be uniform across the traits a breeder cares about. The cultivated tepary pool is a strong starting point for drought improvement: it combines roughly twice the growth rate of wild material with no measurable loss of proportional resilience under water limitation.

Two caveats belong with that result, and both have precedent. The first is that a proportional index is a ratio. Selection on stress-to-control ratios has been known since Rosielle and Hamblin (1981) to favor genotypes with low yield potential rather than genuinely tolerant ones. Thiry *et al*. (2016) restated that bias for modern screening, and Morton *et al*. (2024) observed it directly in a *Solanum pimpinellifolium* panel under salinity, where ratio-based indices overestimated accessions that performed poorly when well watered. That same study found shoot fresh mass under control conditions to explain 73% of the variation in performance under stress, which closely parallels the dominance of cumulative evapotranspiration in our own trait model. The second caveat is transferability. Six accessions re-phenotyped in an independent facility did not reproduce the panel ranking (**Fig. S11**). Our assay also captured early vegetative drought rather than the reproductive-stage stress in which field yield is determined, and genotypic variation for combined drought and heat tolerance in tepary has recently been demonstrated across environments at that stage (Barrera *et al*., 2024).

Our most consequential methodological finding is that plant size determined much of what this experiment appeared to show, entering the traits through normalization and ratios. The mechanism is not new. Traits expressed per unit biomass change with plant size even when the underlying process does not, so an allocation ratio is in large part a size statistic (Poorter and Sack, 2012). Image-derived drought responses likewise decompose into a dominant biomass component and much smaller physiological ones (Chen *et al*., 2014). What this panel adds is how far the effect can carry. Evapotranspiration per unit fresh weight is a standard water-use index, and it produced the single highest association in this study,−log₁₀ *P* = 9.24 at Chr7:14,196,578 under full irrigation. Size adjustment removed that association entirely, across the whole region rather than only at its peak, and left no suprathreshold SNP anywhere in the genome under either treatment (**Supplementary Results S4**). Four of the six loci nominated under full irrigation are led by this trait, and we report them as size loci rather than as water-use loci.

None of this is peculiar to tepary bean. It is a general hazard for phenotyping platforms, in which size, growth stage and position interact to confound exactly the comparisons the platform was built to make (Poorter *et al*., 2023), and it is sharpest in diversity panels whose germplasm classes differ several-fold in size — which is to say most panels containing both wild and improved material. We therefore recommend reporting both the normalized trait and its size-adjusted residual, and treating any locus that does not survive the second as a size locus until shown otherwise. That recommendation cuts against our own result: this panel contains variation in how large plants get and, so far as we can detect, none in how much water they use for a given size.

That the loci detected under water limitation were not those detected under full irrigation is, by contrast, the expected outcome of genotype-by-environment interaction, whether through conditional neutrality or antagonistic pleiotropy (Des Marais *et al*., 2013). Studies that scan both regimes report the same pattern across species. In maize, only one of 27 quantitative trait loci was shared between well-watered and water-deficit conditions (Sarkar *et al*., 2023), biparental populations gave 93 loci under full irrigation against 41 under stress (Ndlovu *et al*., 2024), and plasticity loci cover genomic regions that the regime-specific scans miss entirely (Djabali *et al*., 2023; Li *et al*., 2018). Bread wheat gives 82 associations under deficit and 72 under irrigation with little overlap (Govta *et al*., 2022), and in barley the candidate alleles change the direction of their effect between regimes, which is antagonistic pleiotropy in the strict sense (Arenas and Cortés, 2026). The separation held at every level of evidence, so it is not an artifact of where a threshold falls (**Fig. 5A**). Selection for growth under irrigation is therefore unlikely to move the alleles that matter under drought.

Resolution, not detection, is what limited this study. Mating system governs the physical extent of linkage disequilibrium, and in a self-pollinating species recombination seldom mixes lineages, so haplotypes persist over distances that in an outcrosser would long since have been broken up (Flint-Garcia *et al*., 2003). In our panel uncorrected *r*Z was still declining at 1 Mb, and most associated regions therefore contain tens to hundreds of genes. A single genome-wide decay distance is in any case a poor basis for drawing intervals, because in common bean linkage disequilibrium is distributed very unevenly along chromosomes (Blair *et al*., 2018). We replaced it with a per-locus interval measured on the analyzed genotypes after removing population structure. That measurement is what made gene nomination possible at three of the nine loci and impossible at the rest. It narrowed the chromosome 9 cytokinin oxidase interval to 2.5 kb and the chromosome 11 intervals to 26 and 71 kb, it confirmed that the chromosome 8 block cannot be resolved at all, and it exposed how much of the apparent linkage was ancestry rather than physical proximity — 7 Mb of it on chromosome 8. This is the limitation on association mapping in structured, slowly recombining panels that Korte and Farlow (2013) set out, in a form that can be measured rather than assumed. Two further constraints belong with the mapping. Sequencing depth was moderate, so rare alleles are under-detected. Our permutation scheme also shuffled the phenotype against a fixed kinship matrix, which breaks the correspondence between phenotype and relatedness and can give thresholds that are too permissive (Abney, 2015; John *et al*., 2024b). That is one reason we prefer the Bonferroni threshold here.

The chromosome 8 block illustrates a tension worth stating plainly. It reached−log₁₀ *P* = 8.99 at Chr8:40,059,877, the strongest drought association in the study, and did so for nine measurements. Its allele, however, was at a frequency of 0.91–1.00 in wild accessions and 0.06–0.09 in cultivated ones, germplasm class accounted for up to 86% of the variance in genotype, and the association survived in neither group tested alone. It is a domestication contrast expressed under stress rather than a drought-response allele, and across 3.5 Mb of gene-dense, variant-dense sequence it resolves to no gene. Strength of association and interpretability do not coincide, and we have let the second decide which loci this work follows while still reporting the first.

That result is more interesting than a gene would have been. The alleles that best predicted how much digital biomass a tepary plant accumulated under water deficit are the alleles that most sharply distinguish wild tepary from cultivated forms, and they acted throughout the drought rather than being switched on by it. A substantial part of the genetic variation for performance under water deficit therefore sits in haplotypes that domestication has already sorted. That is at once an argument for returning to wild germplasm and a warning that doing so means moving megabases rather than genes. Distinguishing a large introgression from an inversion or a sweep will require long-read assemblies of contrasting accessions or a recombinant population, not a larger panel.

The chromosome 9 locus points at a specific chemistry, though only at family level. Its candidates include a tandem pair of copper radical oxidases, a family that oxidizes aldehyde substrates including glyoxal and methylglyoxal and generates hydrogen peroxide in the process (Daou and Faulds, 2017). Domain content is a weak guide to substrate in this family, where structure–function work has repeatedly found homologs that barely turn over the substrate their annotation names (Yin *et al*., 2015), so the assignment rests on a family-level expectation rather than a predicted activity.

## Conclusion

We phenotyped a diversity panel of 363 tepary bean accessions through a fourteen-day drought under gravimetric control, scanned 104 measurements for association, and resolved the result to nine independent loci. The strongest is a 3.5 Mb haplotype on chromosome 8, which predicted digital biomass on nine consecutive days of water deficit and separates wild from cultivated accessions almost completely. It is a domestication contrast expressed under stress, and at this resolution it names no gene. The cleanest locus lies on chromosome 11: it is the only one segregating in both wild and cultivated germplasm, it is almost free of population structure, and it localizes to 26 kb, though it points at a receptor kinase and a PAT1-subfamily GRAS factor with no published role in water relations. Between these, three loci yield candidates in which position, linkage interval, expression in droughted tissue and published mechanism agree: a cytokinin oxidase overlapping the lead for chlorophyll index under drought within 2.5 kb, a CIPK1-clade kinase in a module that sets abscisic-acid receptor sensitivity, and LHCB5, which is required for the stomatal response to abscisic acid.

What remains is a question of experimental design rather than of sample size. We measured these associations on shoots, while the mechanisms they implicate — cytokinin catabolism, abscisic-acid receptor sensitivity and stomatal conductance — act through tissues this experiment did not sample. Resolving them will call for expression mapping in those tissues, and for recombinant populations in which the chromosome 8 haplotype can be broken apart and the chromosome 11 interval narrowed to a gene. Tepary bean is already the drought-tolerant member of its genus. This panel shows where in its genome that tolerance is written, and how much of it was sorted on the way to the field.

## Supporting information

Supplementary Results

## Acknowledgements

We thank the staff of the Boyce Thompson Institute Plant Growth Facility — Julie Bell, Tara McCrudden, Philip Ricci and Tim McCullough — for their help in growing the tepary bean panel and in optimizing growth conditions for the stress assays, and Delanie Sickler and Megan Truesdail for coordinating the Boyce Thompson Institute REU program through which A.V. contributed to this work. We thank the USDA National Plant Germplasm System (GRIN) and the International Center for Tropical Agriculture (CIAT) for maintaining and providing the accessions from which the Tepary Diversity Panel was assembled, and Novogene Corporation for whole-genome sequencing.

## Author contributions

AS and LY: investigation, formal analysis, data curation, writing — original draft. RN: investigation (growth habit scoring). AV: investigation. ADLN: methodology, supervision. DP and TP: resources, germplasm curation, funding acquisition, writing — review and editing. MMJ: conceptualization, methodology, formal analysis, supervision, project administration, funding acquisition, writing — original draft, writing — review and editing.

## Conflict of interest

No conflict of interest declared.

## Funding

This work was supported by the United States Department of Agriculture National Institute of Food and Agriculture [award 2022-67013-36212, “Improving Phenomic and Genomic Resources for Resilient relative of Common Bean – Tepary Bean – *Phaseolus acutifolius*”]. Additional support was provided by Boyce Thompson Institute start-up funds to M.M.J.

## Data availability

Raw sequencing reads are deposited at NCBI under BioProject PRJNA1119990. The genome-wide SNP calls — all 6,912,813 called biallelic SNPs across 363 accessions, together with the 6,140,834-SNP filtered set on which the association scans were run, as a bgzipped VCF with index and as the PLINK binary set on which the association scans were run — together with the phenotype matrix, the phenocage images and image-derived canopy traits, association summary statistics, confocal image stacks and the analysis scripts supporting the findings of this study are deposited at Dryad under https://doi.org/10.5061/dryad.9w0vt4bwb Per-SNP summary statistics for the EMMAX replication are included in the deposition alongside the GEMMA scans.

## Notes

https://doi.org/10.5061/dryad.9w0vt4bwb

https://www.ncbi.nlm.nih.gov/bioproject/PRJNA1119990

https://github.com/Leon-Yu0320/Tepary-Genomic-Studies

### Competing Interest Statement

The authors have declared no competing interest.

