## Supplementary Results for "Whole-genome resequencing and indoor phenotyping resolve the genetic architecture of drought stress response in tepary bean"

This file holds the analyses that establish that the associations reported in the main text are properties of the data rather than of the analysis. None of them changes a conclusion; all of them are the reason the conclusions can be stated.

---

### S1. Below the significance threshold the architecture is broadly polygenic

The eight regions reported in the main text are what survives a family-wise threshold, not what the scan contains, and the difference matters for how strongly any one region should be read. Pooling the twenty-one rank-normalized measurements and merging peaks whose lead variants lie within 250 kb gives 307 independent loci reaching $-\log_{10} P = 5$; 96 of these reach 6, and 9 clear the permutation threshold (**Fig. 5**; the complete peak table is included in the data deposition). The threshold therefore sets aside about 97% of the loci the scan nominates. That is the expected picture for a panel of this size and a trait of this kind, and it is why the eight regions are treated as the loci this design can resolve rather than as the loci that exist.

Two patterns in the fuller distribution bear on the main text. Loci are not spread evenly across the genome: chromosomes 8 and 9 carry 38 and 37 loci at $-\log_{10} P \geq 5$ and 19 and 15 at ≥ 6, more than any other chromosome and more than ten times the three loci on chromosome 5 (**Fig. 5**). The concentration of signal on chromosomes 8 and 9 is therefore not an artifact of where the threshold happens to fall. And allele frequency and association strength are unrelated across these loci ($r = -0.06$ between minor allele frequency and $-\log_{10} P$), so the strongest regions are not systematically the ones resting on common alleles: four of the sixteen suprathreshold trait–region combinations have lead variants below a minor allele frequency of 0.15, and the chromosome 8 locus at 54.36 Mb rests on one of 0.082, carried by roughly thirty accessions. The two regions followed in the main text are the exception, with leads at 0.26 (chromosome 9) and 0.37–0.39 (the chromosome 8 block). Recurrence across traits and a common lead allele coincide at those two loci and at neither of the low-frequency single-trait regions, which are not interpreted further.

### S2. The associated regions survive a change of software, and the *AKR4C9* array does not improve

Repeating the scan in EMMAX changes the model in four ways at once — software, kinship matrix, the variants entering that kinship, and the covariates — so it is a demanding test of whether the regions reported are properties of the data or of the analysis. The replication covers all twenty-one rank-normalized measurements (Methods; **Fig. S3**; **Table S8**).

Every region is recovered, with its treatment specificity intact, and EMMAX is systematically the more conservative of the two. Across the fifty-two trait–region combinations that exceed $-\log_{10} P = 5$ in both scans, EMMAX returns the lower value in forty, with a median difference of −1.09 (**Fig. S3A**). Under rank-inverse-normal transformation with the top five principal components as covariates, the chromosome 9 locus reaches 7.16 for digital biomass on day 11 against 8.36 in GEMMA, at the same lead base pair, Chr9:15,146,893; it reaches 2.6 to 7.2 across the twelve drought traits and 2.2 to 3.6 across the nine control traits. The chromosome 8 block behaves the same way but more weakly (6.18 against 8.34), and the control-specific region at Chr8:54,361,120 reproduces most closely of all, at 7.76 against 8.34 for water content under control, with every drought trait below 1.7.

One locus does clear the permutation threshold under EMMAX, and it is worth saying plainly why it is not reported as a finding. A single variant at Chr8:6,611,933 reaches 7.86 for growth rate under control, but no other variant within 250 kb exceeds 1.9, and GEMMA assigns the variant's immediate neighbor a *P* of 0.37. A second EMMAX-only variant, at Chr10:30,680,651, behaves identically (7.68, with nothing above 1.2 beside it). Neither is a rare-variant artifact — the first has a minor allele frequency of 0.13 and a clean genotype distribution — but an association tagged by linkage disequilibrium carries its neighbors with it, and these do not: for comparison, the chromosome 9 lead is flanked by 85 variants above 4 and 265 above 3 in the same scan (**Fig. S3C**). Both are treated as artifacts of holding the variance ratio fixed across variants, which is the one modeling choice that distinguishes EMMAX from GEMMA, and both fall in control traits, which are the traits most confounded with germplasm class. On the same diagnostic, one region that *is* reported — Chr8:54,361,120 — also rests on a lead with little neighboring support; it is retained because both programs find it independently, at 8.34 and 7.76, which neither EMMAX-only variant achieves.

Two further results bear on the interpretation. First, the tandem *AKR4C9* array does not improve under any arm: its best value is 5.81, against 6.05 in the GEMMA scan, and it remains drought-specific — 4.1 to 5.8 across the twelve drought traits against 2.1 to 3.3 under control. The array's weak signal is therefore a property of the data rather than of the association model, which is the more useful of the two possibilities to have established. Second, one region moves the other way. Chromosome 5 at 4.26 Mb is the only region in the comparison where EMMAX exceeds GEMMA, and with the full trait set it does so for every one of the ten drought traits that reach 5 in both scans (+0.04 to +0.72, against a global tendency of −1.09), peaking at 7.50 for growth rate under drought. It is drought-exclusive, with every control trait below 1.5. It clears no threshold in either scan and no gene is nominated, but a region that rises against a uniform conservative shift, in one treatment only and across ten measurements, is the one new lead these analyses produced.

### S3. The untransformed scans are a diagnostic, not a result

Association on untransformed traits behaves as the transformation diagnostic predicts. Growth rate under control conditions, untransformed, yields thirty suprathreshold variants across eight chromosomes, half at minor allele frequency below 0.10; rank-normalized, the same trait yields none anywhere in the genome. Growth rate under drought yields thirty-five untransformed and one rank-normalized. Genomic inflation is 1.02 and 1.05 for these two scans, so this is a heavy phenotype tail producing a few very large test statistics at rare alleles, not global inflation.

In the untransformed EMMAX arms — run on the original nine traits — the proportion of variants below $P = 1 \times 10^{-4}$ reaches 3.7 times its expectation for evapotranspiration per fresh weight, whose strongest signal then falls on chromosome 11 at a position no other trait and no GEMMA scan nominates; under transformation that proportion falls to 0.34 times expectation and the same trait's strongest signal moves into the chromosome 8 block. The untransformed scans are reported in **Table S8** as a diagnostic, not as findings.

### S4. No locus is associated with water use independently of plant size

Because evapotranspiration per fresh weight is largely inverse size (main text), both it and the size-adjusted residual were scanned, to separate a locus acting on water use from a locus acting on size. Evapotranspiration per fresh weight reaches 7.26 at the chromosome 9 locus — its highest value for any trait in this study — and 8.09 in the chromosome 8 block. Both disappear under size adjustment: the residual of log evapotranspiration on log fresh weight gives 3.97 at chromosome 9 and 3.53 in the chromosome 8 block, and has no suprathreshold variant anywhere in the genome under either treatment (genome-wide best 6.97 under drought, 5.66 under control). In the chromosome 8 block the allele that lowers growth rate raises evapotranspiration per fresh weight — opposite signs at the same variants, which is what a size locus looks like when size sits in the denominator.

There is therefore no genetic evidence in this panel for a locus acting on water use independently of plant size.
