## Supplementary Figures and Table legends for "Whole-genome resequencing and indoor phenotyping resolve the genetic architecture of drought stress response in tepary bean"

#### Contents

**Supplementary Figures S1–S12** — one per page, each with its legend beneath.

**Supplementary Tables S1–S16** — legends listed at the end of this document. The tables themselves are provided as a separate spreadsheet.

### Supplementary Figures

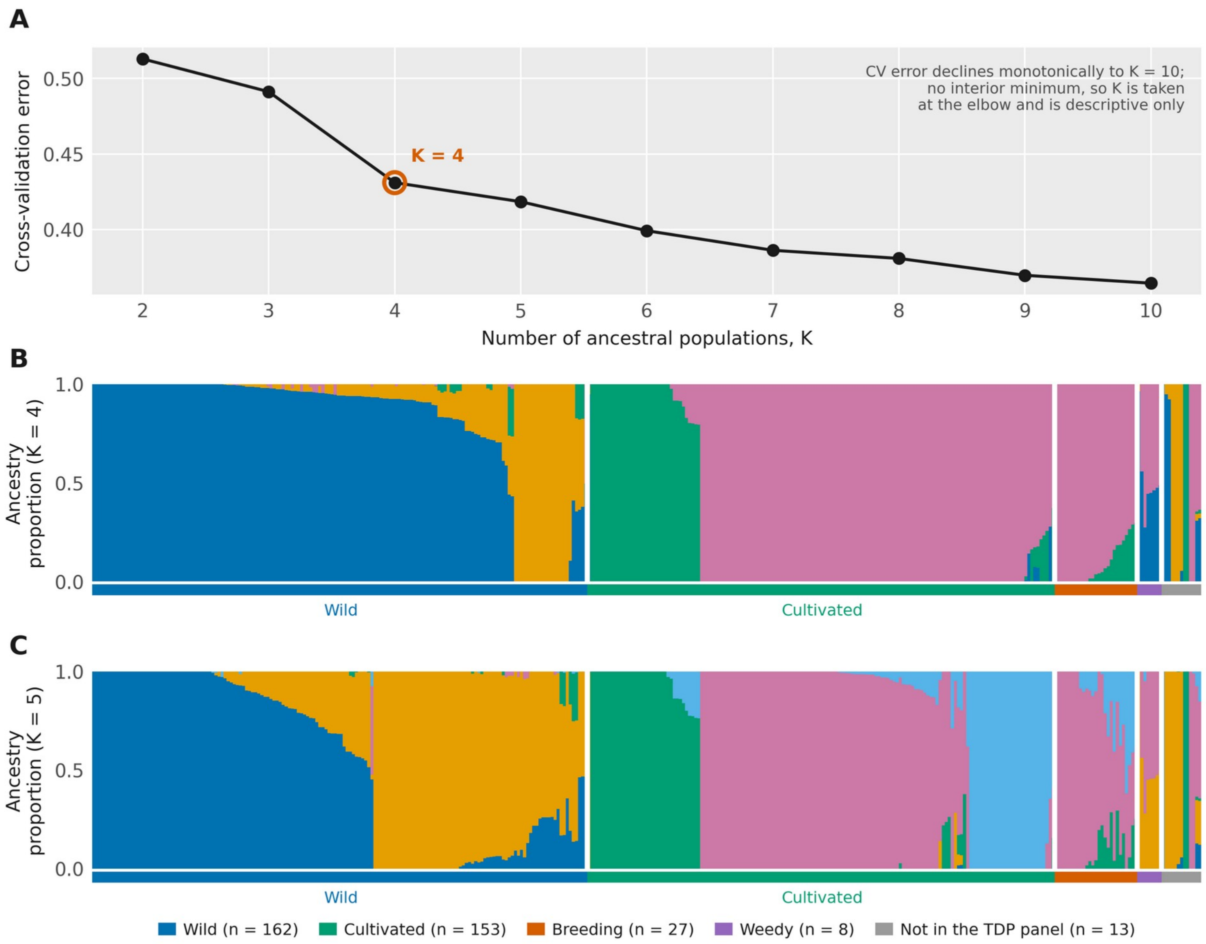


**Figure S1.** Cross-validation does not identify a single best K, but the cultivated pool resolves as one ancestry component at every K tested while wild germplasm splits into several. **(A)** ADMIXTURE cross-validation error for K = 2 to 10, estimated on 65,596 variants (the linkage-disequilibrium-pruned set of **Fig. 1C, D** further filtered to variants missing in fewer than 10% of accessions). The error falls monotonically across the whole range and has no interior minimum, so cross-validation does not identify K in this panel — the expected outcome when structure is continuous rather than a small number of discrete populations. The largest single reduction is at K = 4 (0.431, from 0.491 at K = 3), after which the curve flattens; K = 4 is therefore used as a descriptive summary and not as an estimate of the number of ancestral populations. Bornowski *et al.* (2023) report K = 6 for their panel, which falls in the flat part of this curve and is not in conflict with it. **(B)** Ancestry proportions at K = 4, one stacked bar per accession, ordered by germplasm class and then by dominant component; the ribbon beneath each panel gives germplasm class. The cultivated pool is captured almost entirely by a single component (90.2% of cultivated accessions assigned to it above 0.9; mean maximum ancestry 0.977), and breeding lines share that component. Wild accessions divide between two components (mean maximum ancestry 0.921). The eight weedy accessions are the only group that is consistently admixed (mean maximum ancestry 0.616; one of eight above 0.9), consistent with weedy tepary bean carrying both crop and wild ancestry. Thirteen sequenced accessions have no passport record in the TDP panel and are shown separately. **(C)** The same accessions at K = 5, shown for comparison: the additional component subdivides the cultivated pool and leaves the wild–cultivated split unchanged.

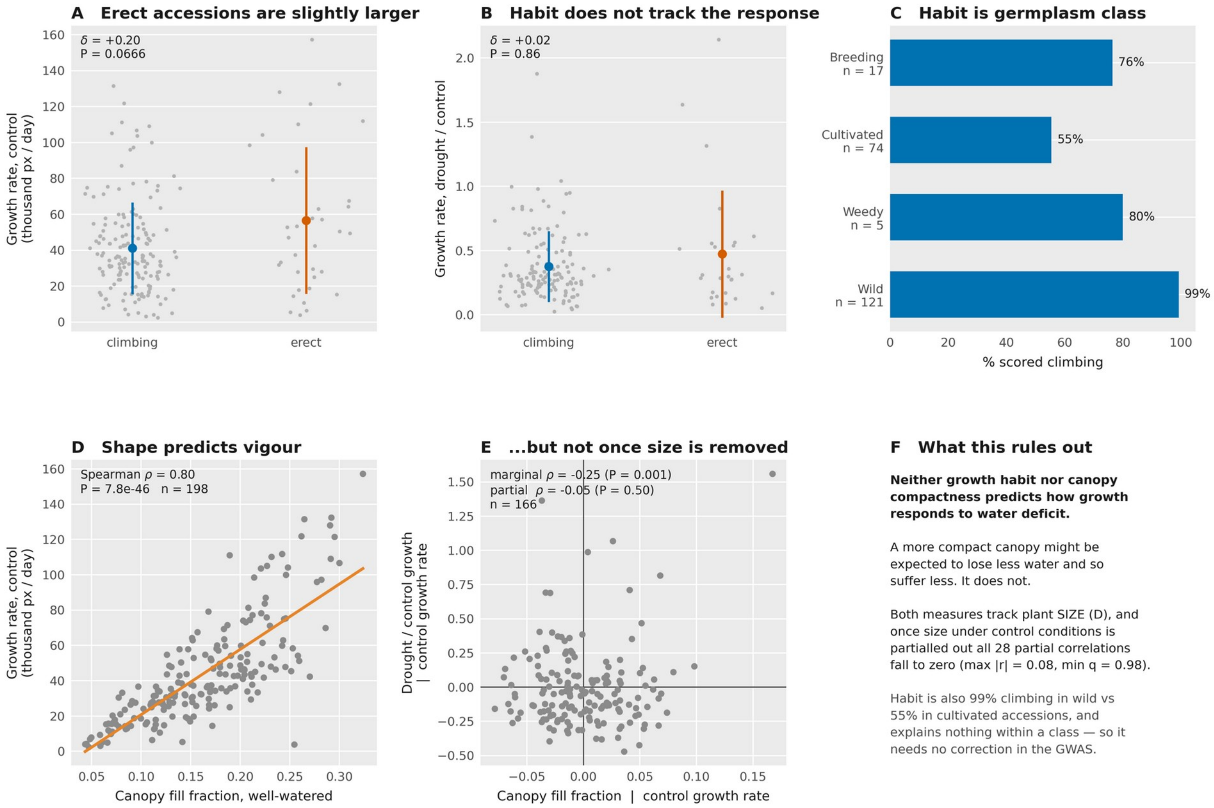


**Figure S2.** Neither growth habit nor canopy shape predicts the drought response. **(A)** Growth rate under control conditions by scored growth habit; erect accessions are somewhat larger, though not significantly so in the 229 accessions with both a habit call and panel phenotypes (Cliff's δ = +0.16, *P* = 0.12). **(B)** The drought/control growth-rate ratio by habit: no difference (δ = +0.05, *P* = 0.68). **(C)** Proportion of accessions scored climbing within each germplasm class — habit is very largely wild-versus-cultivated status. **(D)** Canopy fill fraction measured on a single well-watered phenocage image against growth rate under control conditions in the separate panel experiment (Spearman ρ = 0.82, $P = 1 \times 10^{-51}$, *n* = 206): canopy shape is a strong cross-experiment proxy for vigor. **(E)** The same two variables against the drought response, each with control-condition growth rate partialed out. The marginal correlation (ρ = −0.28, $P = 3 \times 10^{-4}$) collapses to nothing once size is removed (partial ρ = −0.08, *P* = 0.28). **(F)** Summary. Ratios outside the interval (0, 3] were excluded as uninterpretable, as were the negative growth rates described in the Methods.

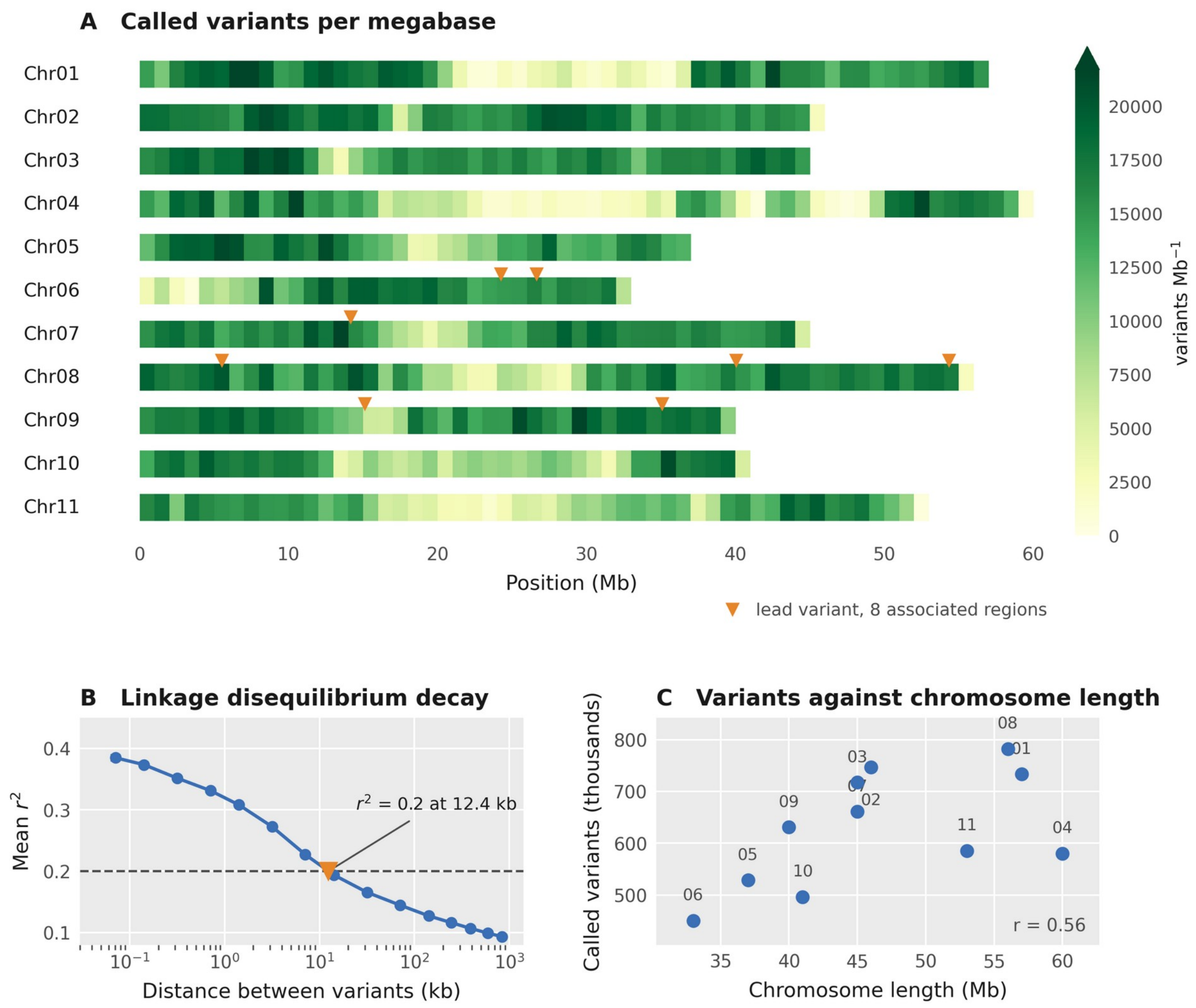


**Figure S3.** Variant density is uneven across the genome and linkage disequilibrium decays slowly. Mean $r^2$ against physical distance between variants, in logarithmically spaced bins, for 111,305 randomly thinned variants across the 226 phenotyped accessions (28.2 million pairs within 1 Mb). Error bars, standard error of the bin mean. Dashed line, $r^2 = 0.2$; the marker gives the interpolated crossing at 12.4 kb. This genome-wide average is reported for comparability with other panels; the candidate intervals used here were measured per locus after correcting for population structure (**Table S12**). $r^2$ is still declining in the outermost bin: no plateau is reached within 1 Mb.

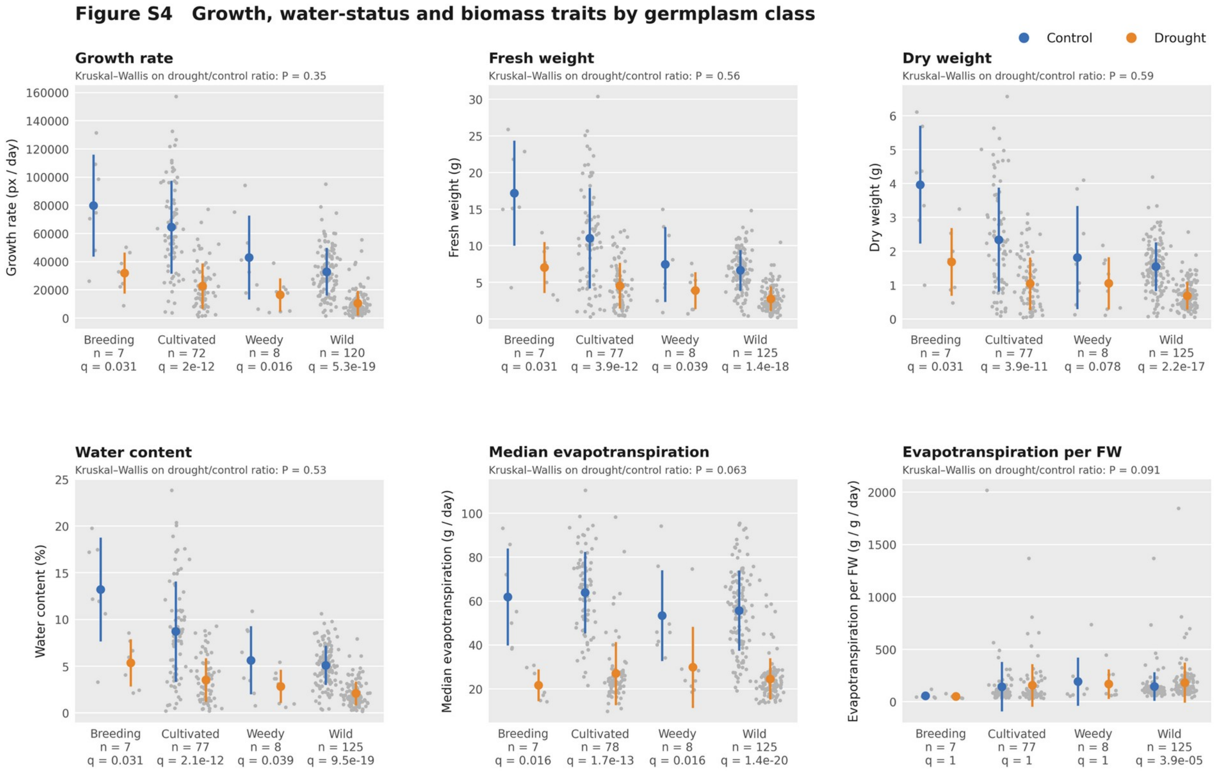


**Figure S4.** Cultivated and breeding lines are larger than wild accessions under both watering regimes. Growth rate, fresh weight, dry weight, tissue water content, median evapotranspiration and evapotranspiration per unit fresh weight, under control (blue) and drought (orange) within each germplasm class. Gray points, individual accessions; colored point and bar, mean ± SD. *q*, paired Wilcoxon signed-rank test within class, Holm-adjusted across the four classes; *n*, accessions with a value under both conditions. Above each panel, the Kruskal-Wallis test of whether the drought/control ratio differs between classes. Impossible values were excluded (Methods).

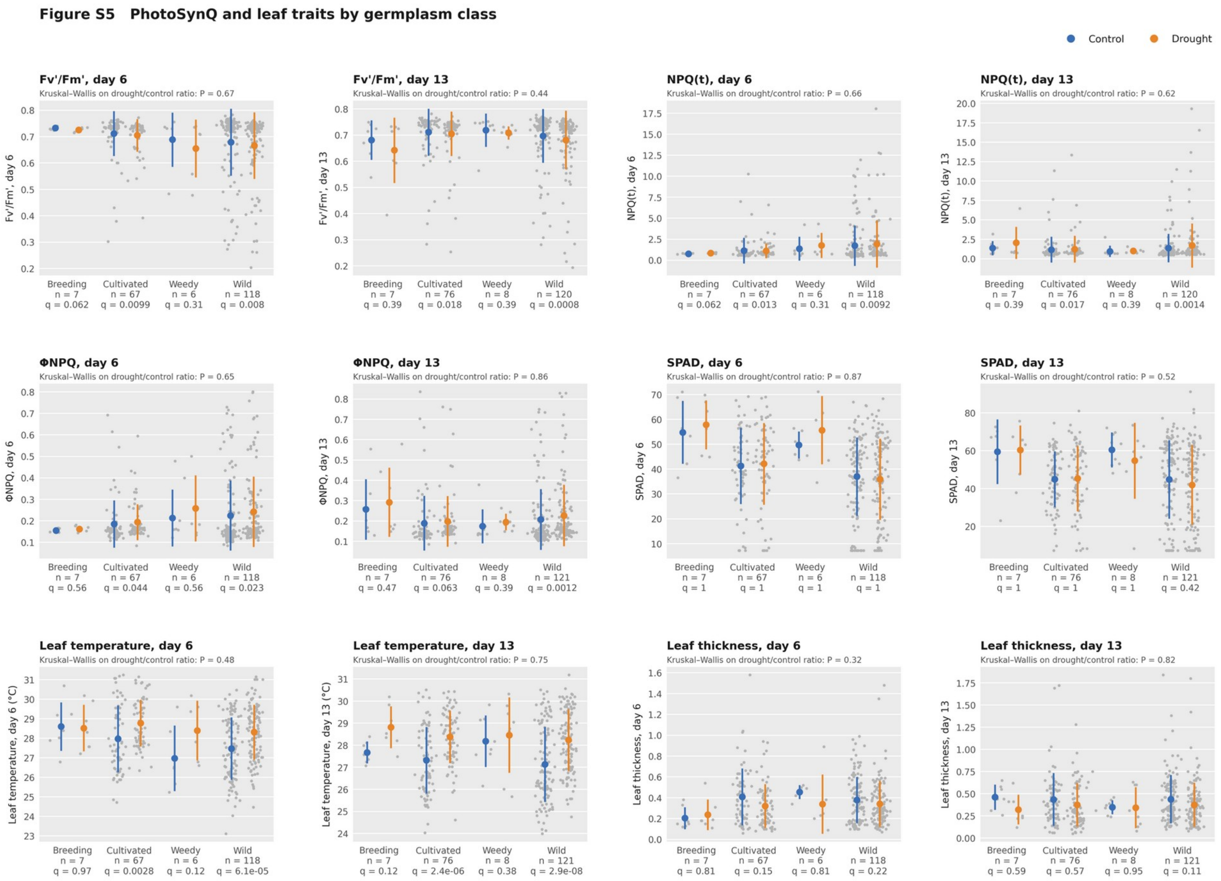


**Figure S5.** Photosynthetic parameters differ little between germplasm classes under either regime. Fv′/Fm′, NPQ(t), ΦNPQ, SPAD, leaf temperature and leaf thickness, on days 6 and 13 after stress imposition. Conventions as in Figure S4.

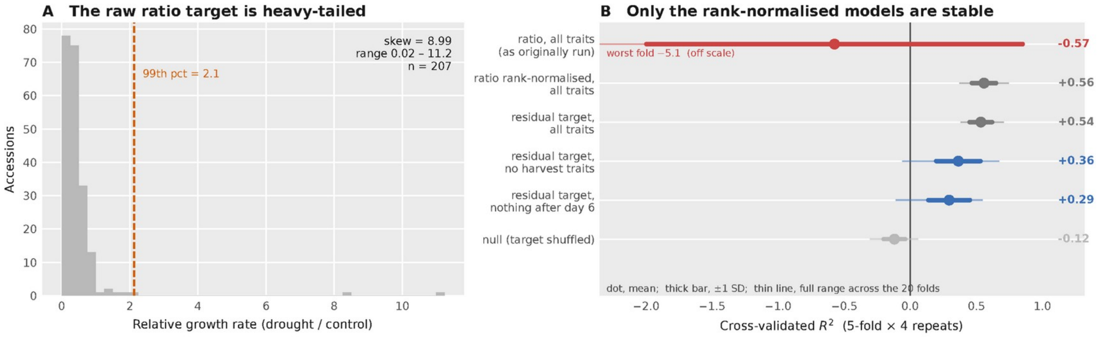


**Figure S6.** Why the raw drought/control ratio cannot be used as a regression target. **(A)** Distribution of the relative growth rate (drought/control) across the 207 accessions analyzed; dashed line, 99th percentile. **(B)** Cross-validated $R^2$ for six model specifications; dot, mean over 20 fits (5 folds × 4 repeats); thick bar, ±1 SD; thin line, full range across folds. The model as originally specified (red) has a mean $R^2$ below zero and a fold range from −5.1 to +0.7. These panels were Figure 4A, B in an earlier version and retain British spelling in their axis labels.

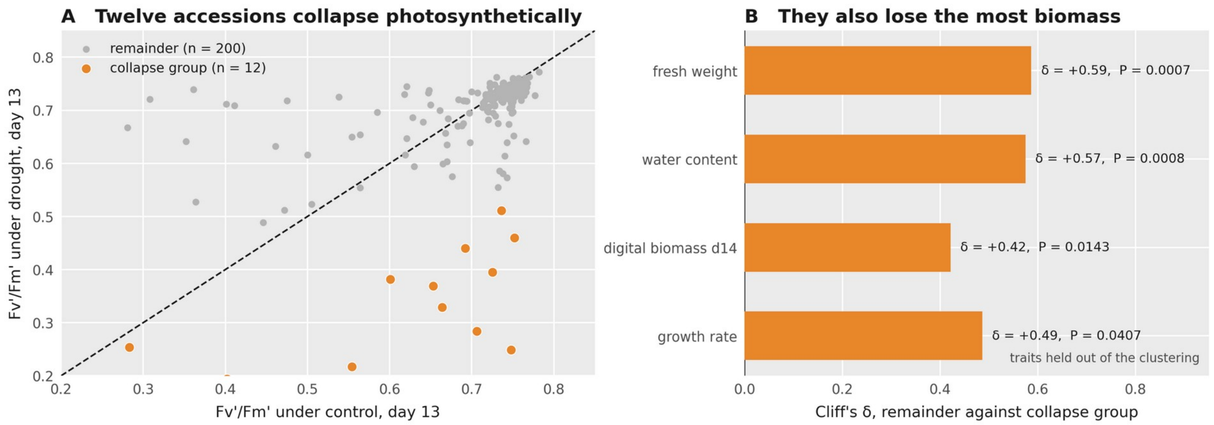


**Figure S7.** Twelve accessions shut down photosynthesis under drought and maintain growth poorly. **(A)** Fv′/Fm′ on day 13 under drought against the same accession's control value; orange, the twelve accessions separated by clustering on the photosynthetic response alone (silhouette 0.75). **(B)** Cliff's δ between the two groups for four traits held out of the clustering; the collapse group loses more fresh weight, water content, digital biomass and growth. The grouping is independent of germplasm class ($\chi^2$, $P = 0.55$) and is not a watering artifact — these accessions transpired less from day one and started at the same size. With twelve accessions and no independent experiment this is an observation requiring replication.

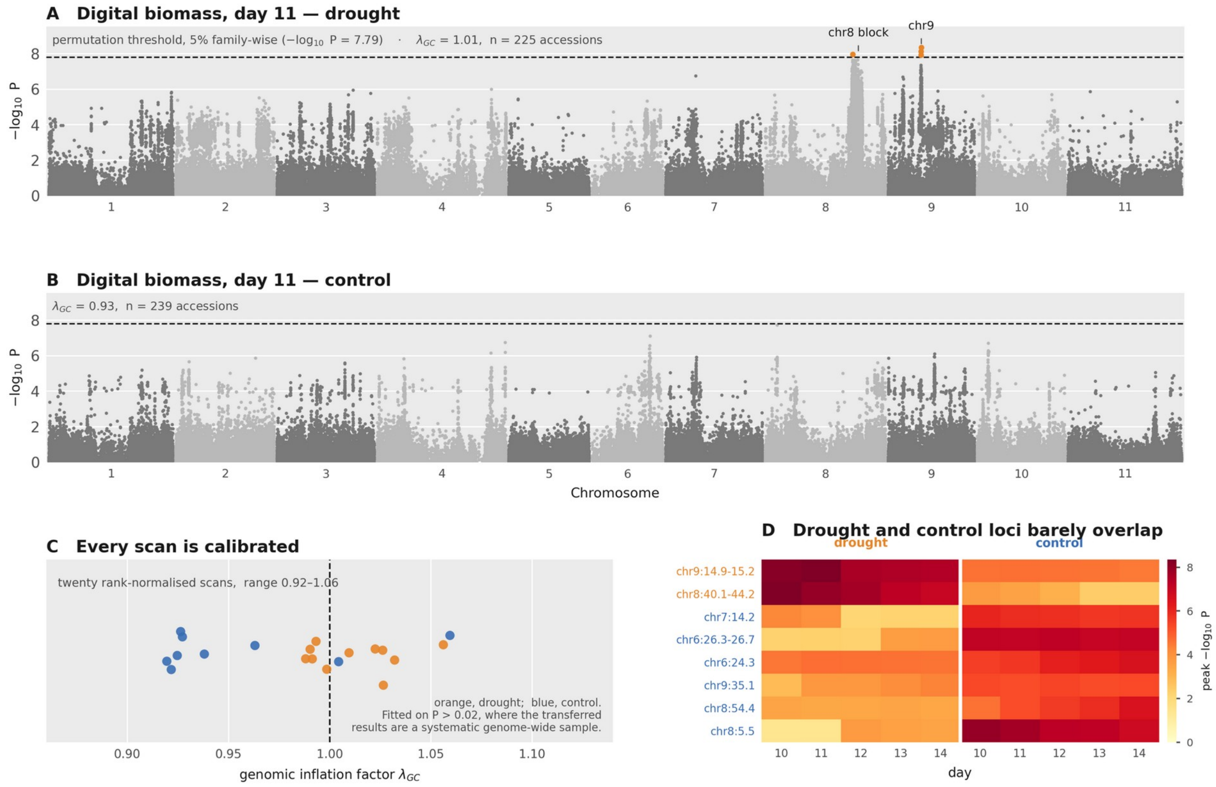


Association is shown for the twenty rank-normalised trait–treatment combinations, which are the primary scan of this study; the untransformed scans are reported only as a diagnostic (Results). Each panel is the likelihood-ratio test from a linear mixed model in GEMMA over 5.80–5.89 M variants with a centred relatedness matrix. The threshold is the fifth percentile of the genome-wide minimum P across 100 phenotype permutations, $-\log_{10}$ P = 7.79 (Methods); it replaces the Bonferroni-style value used in earlier versions of this analysis. Twenty-three variants pass it, in eight loci — two specific to drought, six to control. The five suprathreshold stretches on chromosome 8 between 40.06 and 44.23 Mb are one continuous block and are counted once.

**Figure S8.** Diagnostics for the digital biomass scans. **(A, B)** The Manhattan plots reproduced as **Fig. 6A, B**. **(C)** Genomic inflation factor for each of the twenty-one rank-normalized scans, orange for drought and blue for control, fitted on the part of each scan above $P = 0.02$ where the thinned results are a systematic genome-wide sample; all twenty lie between 0.92 and 1.06. **(D)** Peak $-\log_{10} P$ within each of the nine loci, on days 10–14 under each condition; row labels are colored by the treatment the locus belongs to.

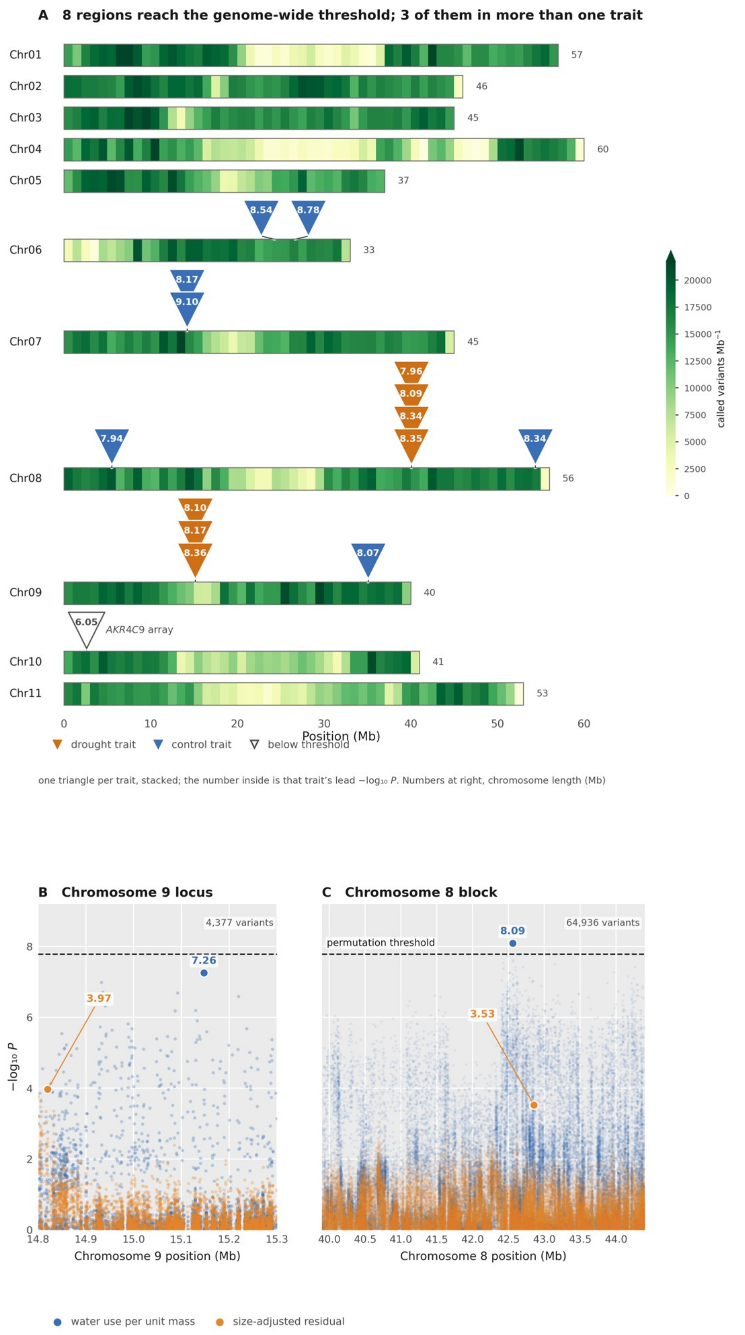


**Figure S9.** Association overview: which traits each region is significant for, and water-use associations before and after adjustment for plant size. **(A)** The eight suprathreshold regions on a variant-density backbone; superseded by **Fig. 5**, retained here because panels B and C share its coordinate frame. **(B)** Which traits each region is significant for. Every region is significant for traits measured under one watering regime only: the drought and well-watered scans share no region. **(C)** Water-use associations before and after adjustment for plant size. Blue, evapotranspiration per unit fresh weight; orange, the residual of log evapotranspiration on log fresh weight. The chromosome 9 and chromosome 8 signals fall from 7.26 and 8.09 to 3.97 and 3.53, and the size-adjusted trait has no suprathreshold variant anywhere in the genome under either treatment (**Supplementary Results S4**).

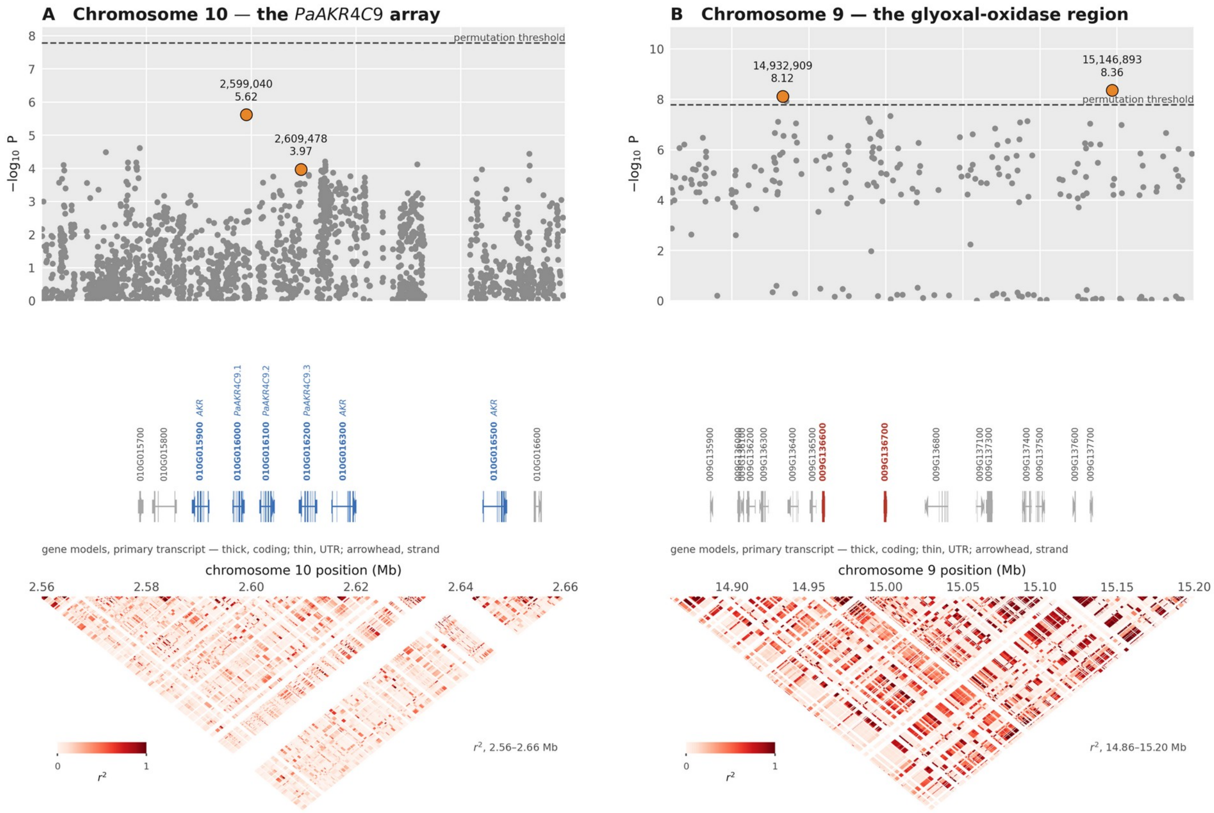


**Figure S10.** The chromosome 9 reactive-carbonyl locus: full gene models and variant consequences. **(A, B)** Gene models across the chromosome 9 copper-radical-oxidase region, reproduced as **Fig. 6C**; models are the primary transcript of each gene in the *P. acutifolius* v1.0 annotation (Moghaddam *et al.*, 2021). **(C)** The haplogroup analysis reproduced as **Fig. 6D**. **(D)** Every variant reaching significance in the region, with its predicted consequence.

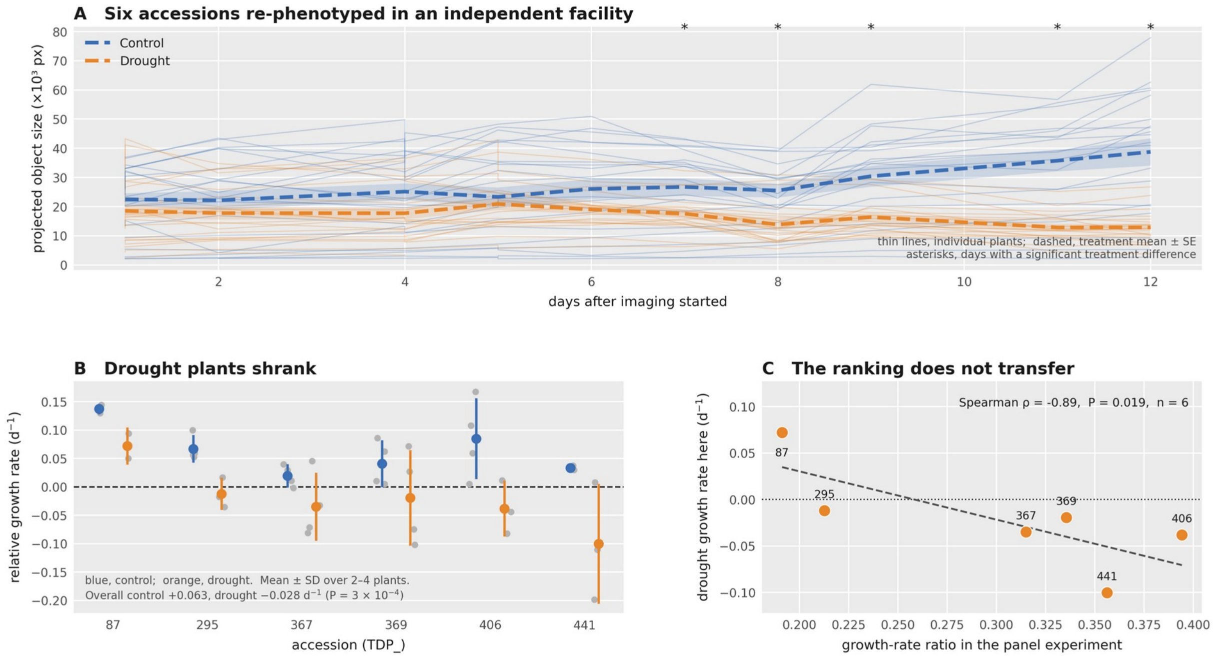


Six accessions were re-grown in the PhenoSight facility under a control and a drought regime, 2–4 plants per accession per treatment, imaged over 16 days; days with fewer than 35 of the 41 trays imaged were excluded, leaving ten timepoints. The drought imposed here was more severe than in the panel experiment: plants lost projected area rather than growing more slowly, so a drought-to-control ratio is undefined and growth rates are compared directly. Chlorophyll fluorescence did not register the stress (Fv/Fm 0.748 control against 0.787 drought, P = 0.16; npq(t) 2.41 against 2.17, P = 0.49), consistent with the weak standing of photosynthetic parameters in the trait model. Accession ranking did not transfer between the two experiments and if anything inverted (C) — the accessions that maintained growth best in the panel did worst here. Both endpoints are size-dependent in opposite directions, which is the confound documented throughout this study rather than a contradiction between experiments; with six accessions and 2–4 replicates this comparison is underpowered and is reported as a caution, not a result.

**Figure S11.** Independent re-phenotyping does not reproduce the panel ranking. **(A)** Projected object size against time for every plant, under control (blue) and drought (orange); thin lines, individual plants; dashed lines, treatment mean ± SE; asterisks mark days on which the treatments differ (Mann–Whitney, $P < 0.05$). Days on which fewer than 35 of the 41 trays were imaged are excluded, leaving ten timepoints. **(B)** Relative growth rate fitted per plant over that window, by accession and treatment; gray points, individual plants; colored point and bar, mean ± SD over 2–4 plants. **(C)** Drought growth rate here against the drought/control growth-rate ratio recorded for the same accessions in the panel experiment; dashed line, ordinary least squares. **(D)** Analysis note. Chlorophyll fluorescence did not register the treatment (Fv′/Fm′ 0.748 control against 0.787 drought, $P = 0.16$; npq(t) 2.41 against 2.17, $P = 0.49$).

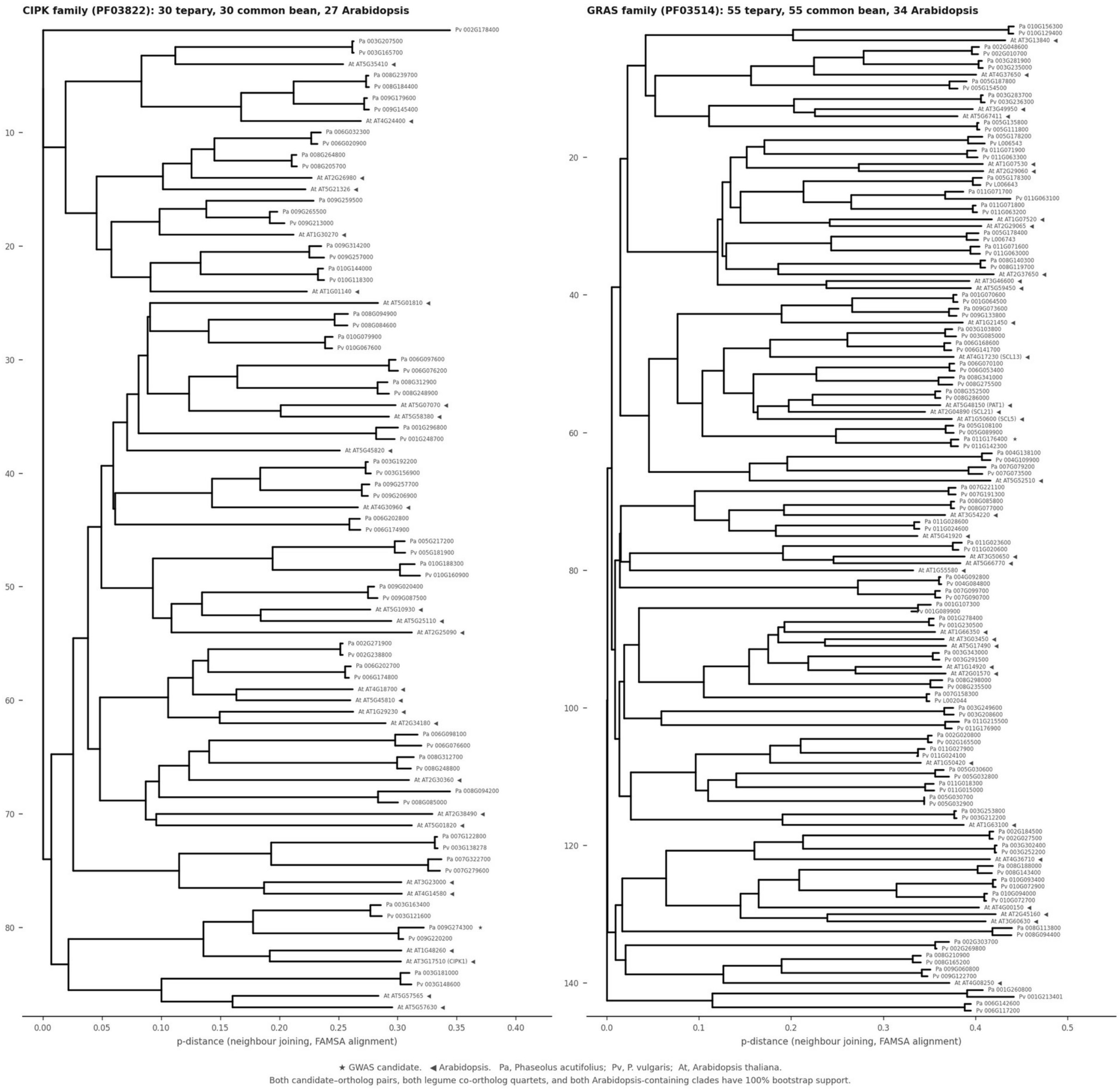


**Figure S12.** Family phylogenies for the CIPK and GRAS candidates. Neighbor-joining trees of the CIPK (PF03822, left; 30 tepary, 30 common bean, 27 Arabidopsis) and GRAS (PF03514, right; 55, 55 and 34) families, aligned with FAMSA and built on p-distance over non-gap positions, midpoint-rooted. Tips are prefixed Pa (*P. acutifolius*), Pv (*P. vulgaris*) and At (*Arabidopsis thaliana*); Arabidopsis tips are marked with a triangle and the genes named in the text are labeled. A star marks the GWAS candidate in each family. Bootstrap support from 200 (CIPK) and 100 (GRAS) column-resampling replicates is 100% for both candidate–ortholog pairs, for both legume co-ortholog quartets, and for the clades uniting the CIPK quartet with CIPK1 and the GRAS quartet with the PAT1 subfamily.

#### Supplementary Table legends

The tables are supplied as a separate spreadsheet; the legends are reproduced here.

**Table S1.** Germplasm information for the tepary bean accessions used in this study. All 363 sequenced accessions are listed: accession identifier, source collection, germplasm class (wild, weedy, cultivated or breeding line), country and state or department of origin, taxonomic assignment, and the source of the sequence data. The 13 accessions taken from Moghaddam *et al.* (2021) are identified there and carry their SRA run identifiers in place of a TDP number.

**Table S2.** Sequencing and read-mapping statistics per accession: total paired reads and bases generated, mean depth, reads mapped to the *P. acutifolius* reference, and mapping rate.

**Table S3.** Variant-calling statistics per accession.

**Table S4.** Genotype-specific phenotypic data from the controlled-environment experiment. One row per accession; columns give growth rate, fresh and dry weight, tissue water content, daily and cumulative evapotranspiration, digital biomass on days 0–14, and the PhotoSynQ parameters recorded on days 6 and 13, each under control and drought conditions. Fourteen of the drought and nine of the control growth-rate values are negative; these are curve-fit artifacts and are excluded from every analysis reported here (Methods).

**Table S5.** Distribution statistics for the digital biomass and growth traits: Shapiro-Wilk test, skewness, kurtosis, mean and median per trait, day and treatment, used to choose the transformation applied before association testing.

**Table S6.** The same for the PhotoSynQ photosynthetic traits.

**Table S7.** Traits taken forward to genome-wide association, their units, the transformation applied to each, and the number of accessions contributing. Evapotranspiration and the harvest masses are in grams; the remaining traits are image- or sensor-derived indices reported in arbitrary units (a.u.). Both the transformed and the rank-inverse-normal form of every trait were tested.

**Table S8.** The 248 annotated genes within 150 kb of the seven regions for which genes are listed, with distance to the region's lead variant, whether the gene falls inside that locus's structure-corrected candidate interval, best Arabidopsis and rice hits, gene description and Pfam domains; 154 carry a functional assignment. Regions contain 20 to 77 genes, so proximity is not evidence of causality. The chromosome 8 block is excluded.

**Table S9.** Repositories holding the raw and processed data supporting this study — sequencing reads, variant calls, the phenotype matrix, phenocage images and derived canopy traits, association summary statistics, and the analysis scripts.

**Table S10.** Growth habit and canopy shape per accession, scored from side-view images across eight phenocage experiments: consensus habit call, mean spindliness score, canopy fill fraction and aspect ratio under each treatment, and the number of images contributing.

**Table S11.** Correlations between canopy architecture and evapotranspiration, before and after partialing out fresh weight, with Benjamini-Hochberg adjusted *P*-values.

**Table S12.** The nine independent loci reaching the Bonferroni threshold ($-\log_{10} P$ = 8.07), one row per merged region. The first two columns give the locus each region belongs to and its category — domestication contrast, drought-specific, or full irrigation, the last subdivided into size loci, the structure-independent chromosome 11 locus and the provisional Chr8:54.36 Mb region. The remaining columns give: chromosome, span, lead variant position and $-\log_{10} P$, minor allele frequency, effect size, call rate and heterozygosity; uncorrected and structure-corrected linkage-disequilibrium intervals at $r^2 \geq 0.6$ and $\geq 0.3$; allele frequency in each germplasm class and the proportion of genotype variance attributable to class; and the association re-fitted without covariates, with five principal components, with class as a covariate, and within the Cultivated and Wild groups separately.

**Table S13.** Quantitative PCR primers for the two chromosome 9 copper radical oxidases, *STH2*, and two reference genes. Every pair was verified by in-silico PCR against the complete *P. acutifolius* transcriptome.

**Table S14.** Every annotated gene inside the structure-corrected candidate interval of each of the nine loci (322 genes), with gene model coordinates, strand, distance to the nearest suprathreshold variant, whether the gene falls inside the $r^2 \geq 0.6$ core or only the outer interval, best Arabidopsis and rice hits, gene description, Pfam domains and Gene Ontology terms.

**Table S15.** Expression of the 37 candidate genes across four resources: mean FPKM in each of twelve tepary tissues and across the leaf heat time course (Moghaddam *et al.*, 2021); the reciprocal-best-hit common bean ortholog and whether the hit was reciprocal; baseMean, $\log_2$ root/leaf ratio and expression percentile in droughted field-grown common bean (Subramani *et al.*, 2023); and drought-versus-control fold changes where the ortholog is differentially expressed in Gregorio Jorge *et al.* (2020) or López *et al.* (2020). A verdict column records how much expression evidence supports each candidate.

**Table S16.** The twelve accessions that shut down photosynthesis under drought. Accession identifier, source accession ID and germplasm class, followed by the day-13 PhotoSynQ parameters used for the clustering — Fv′/Fm′, NPQt, ΦNPQ and SPAD — each under control and under drought and as the residual of the drought value on the control value, and the four traits held out of the clustering (growth rate, fresh weight, water content and digital biomass on day 14) under both regimes. The group was defined by *k*-means clustering on the four photosynthetic residuals alone (silhouette 0.75; **Fig. S7**). Values are per-accession means. *(file: TableS_photosynthesis_collapse_accessions.csv)*
